# ALK R1275Q mutation drives expansion of SCP-like cells during sympathoadrenal commitment and primes neuroblastoma initiation

**DOI:** 10.64898/2026.01.27.701690

**Authors:** Mingzhi Liu, Panagiotis Alkinoos Polychronopoulos, Ana Marin Navarro, Veronica Zubillaga, Huaitao Cheng, Qirong Lin, Tzu-Po Chuang, Jinhye Ryu, Lola Boutin, Yiming Xia, Ann-Sophie Oppelt, Leilei Zhou, Hugo Metzger, Martin Enge, Daniel Bexell, John Inge Johnsen, Oscar C. Bedoya-Reina, Ruth Palmer, Per Kogner, Anna Falk, Margareta Wilhelm

**Author notes:** Correspondence to (M.W.).

## Abstract

Neuroblastoma (NB) is a pediatric malignancy developing in the sympathoadrenal lineage of the neural crest, characterized by clinical heterogeneity ranging from spontaneous regression to poor outcomes. Activating mutations in the receptor tyrosine kinase anaplastic lymphoma kinase (ALK) are frequently observed in both sporadic and familial NB, yet the functional role of ALK in tumor initiation is not fully understood. Using a patient-derived human induced pluripotent stem cell (iPSC) model of sympathoadrenal development, we show that upon sympathoadrenal lineage commitment, ALK R1275Q, the most common hotspot mutation found in familial NB, sustain a proliferative, immature Schwann cell precursor (SCP)-like cell state with elevated ALK signaling and increased susceptibility to MYCN-driven transformation. While ALK-mutant cells alone did not form tumors *in vivo*, they cooperated with MYCN to accelerate tumor initiation, suggesting that ALK R1275Q creates a permissive but insufficient state for transformation. These findings define an ALK-driven cell progenitor-like state that facilitates the initiation of NB during embryonal development.

## Introduction

Neuroblastoma (NB) is a pediatric malignancy that originates from neural crest-derived cells committed to the sympathetic nervous system lineage, and is considered a prototypical developmental cancer due to its emergence from sympathoadrenal (SA) progenitors ^1–4^. NB is the most common extracranial solid tumor in children and accounts for approximately 15% of pediatric cancer-related mortality ^5,6^. NB is clinically heterogeneous, with some tumors in infants regressing spontaneously, others follow a slow and prolonged course, whereas high-risk NB presents as an aggressive disease that remains difficult to cure despite intensive multimodal therapy ^7–9^. This variability emphasizes the need to understand how disruptions in SA lineage development initiate tumorigenesis and how genetic alterations shape disease trajectory.

Among the genetic alterations implicated in NB, activating mutations in the receptor tyrosine kinase ALK (anaplastic lymphoma kinase) is a well-established familial predisposition factor^10–12^. Recurrent hotspot mutations within the kinase domain, including R1275Q, F1174L, and F1245C, confer constitutive kinase activity contributing to both familial and sporadic cases. R1275Q is the most common germline variant and displays incomplete penetrance, suggesting that additional oncogenic events are required for malignant transformation. In contrast, somatic alterations such as F1174L or ALK amplification often show stronger oncogenic potency and are enriched in high-risk tumors^10,11,13–17^. Together, these findings indicate that ALK mutations are play a role in NB pathogenesis, but their tumorigenic potential is influenced by developmental context and cooperating genetic alterations.

Most NB initiation models have relied on genetically engineered mouse models (GEMMs) and xenograft models, which have been valuable for understanding disease biology, but remain limited in recapitulating human-specific developmental trajectories^18–22^. In particular, these systems may be less suited to identify subtle lineage-specific changes that may occur before tumor formation. Pluripotent stem cell (PSC)–based *in vitro* models, including embryonic stem cells (ESCs) and induced pluripotent stem cells (iPSCs), have emerged as powerful complementary alternatives, that allow precise temporal control of differentiation and investigation of early human developmental states^23–25^. These models have been successfully used to study oncogenic alterations in pediatric cancers, including *MYCN* in NB^26^, *PTCH1* mutation in medulloblastoma^24,27^, H3.3K27 mutation in pediatric glioma^28,29^, *NF1* mutation in neurofibroma^30,31^, and *EWS-FLI1* and ETV6-RUNX1 translocations in Ewing’s sarcoma and B acute lymphoblastic leukemia, respectively^32,33^. While PSC-based studies of ALK mutations, including the germline R1275Q mutation or the more aggressive F1174 mutation, have provided insights^34–36^, previous research mainly focused on tumor phenotypes, leaving their broader developmental relevance to be clarified.

To address these gaps, we developed a human iPSC-based model system to study the consequences of ALK R1275Q mutation during normal sympathoadrenal development. Using three different patient-derived iPSC lines carrying the ALK R1275Q mutation, we found that patient-derived cells deviated from normal differentiation by remaining in a proliferative, immature state. Importantly, we identified a distinct SCP-like subpopulation at the stage of SA lineage commitment, expressing both SCP and SA genes, that expanded and was more persistent in patient-derived cells. Functionally, this subpopulation showed elevated ALK activity and increased susceptibility to MYCN-driven transformation, linking this population to proliferative imbalance and tumor initiation risk. Our findings support a model in which ALK R1275Q acts less as a classical oncogenic “switch” and more as a lineage-modifying factor that prolongs a transformation-prone state.

Our study establishes a human iPSC-based developmental model for ALK-driven NB initiation and reveals an SCP-like, transformation-prone state that may present vulnerabilities for combination strategies with ALK inhibition.

## Result

### *In Vitro* Differentiation of Induced Pluripotent Stem Cells into the Sympathoadrenal Lineage via Trunk Neural Crest Cells

To model SA lineage development, we established a stepwise differentiation protocol by combining and modifying previous protocols^37–39^. iPSCs, derived from three healthy individuals (Ctrl7, Ctrl10, and Ctrl14)^40,41^, were directed into trunk neural crest cells (tNCC), sorted for NGFR/p75^NTR+^ cells, followed by differentiation to sympathoadrenal progenitors (SAPs), and subsequently to sympathoadrenal mature cells (SAMs) (Fig. 1a). All iPSC lines were reprogrammed using non-integrative methods, passed pluripotency quality control assays, and were routinely validated by immunostaining for SSEA4, OCT4, and NANOG (Fig. 1a, Extended Data Fig. 1a and d)^40^. Using the established protocol, iPSCs differentiated into tNCCs in 6 days, which we set as day 0 (D0), SAPs were obtained 4 days later (D4), and SAMs after an additional 8 days (D12) (Fig. 1a, left). At each stage, lineage-defining markers were confirmed at both transcriptional and protein levels, including SOX10 and HOXC9 in tNCCs, and PHOX2B, PRPH, and TUBB3 in SA cells (Fig. 1a, right; Extended Data Fig. 2 and Supplementary Fig. 1-5). To further validate positional identity and lineage-specific gene expression bulk transcriptomic analyses were performed at six sequential time points, ranging from D0 (tNCCs) to D12 SA cells (Fig. 1a). HOX family gene expression analysis confirmed trunk-level NCCs (Fig. 1b, Supplementary Data 1). Furthermore, comparison with published datasets from human adrenal medulla revealed that tNCCs expressed markers of NCCs, Schwann cell precursors (SCPs), and bridge-cells (*SOX10*, *PLP1*, *PAX3*, *ERBB3*, *MPZ*), whereas SA-committed cells (*PHOX2B*) expressed sympathoblast (*HAND2*, *PRPH*, *GATA3*) and to a lesser extent chromaffin cell markers (*CHGA*, *PENK*) (Fig. 1c)^42,43^.

**Figure 1.**
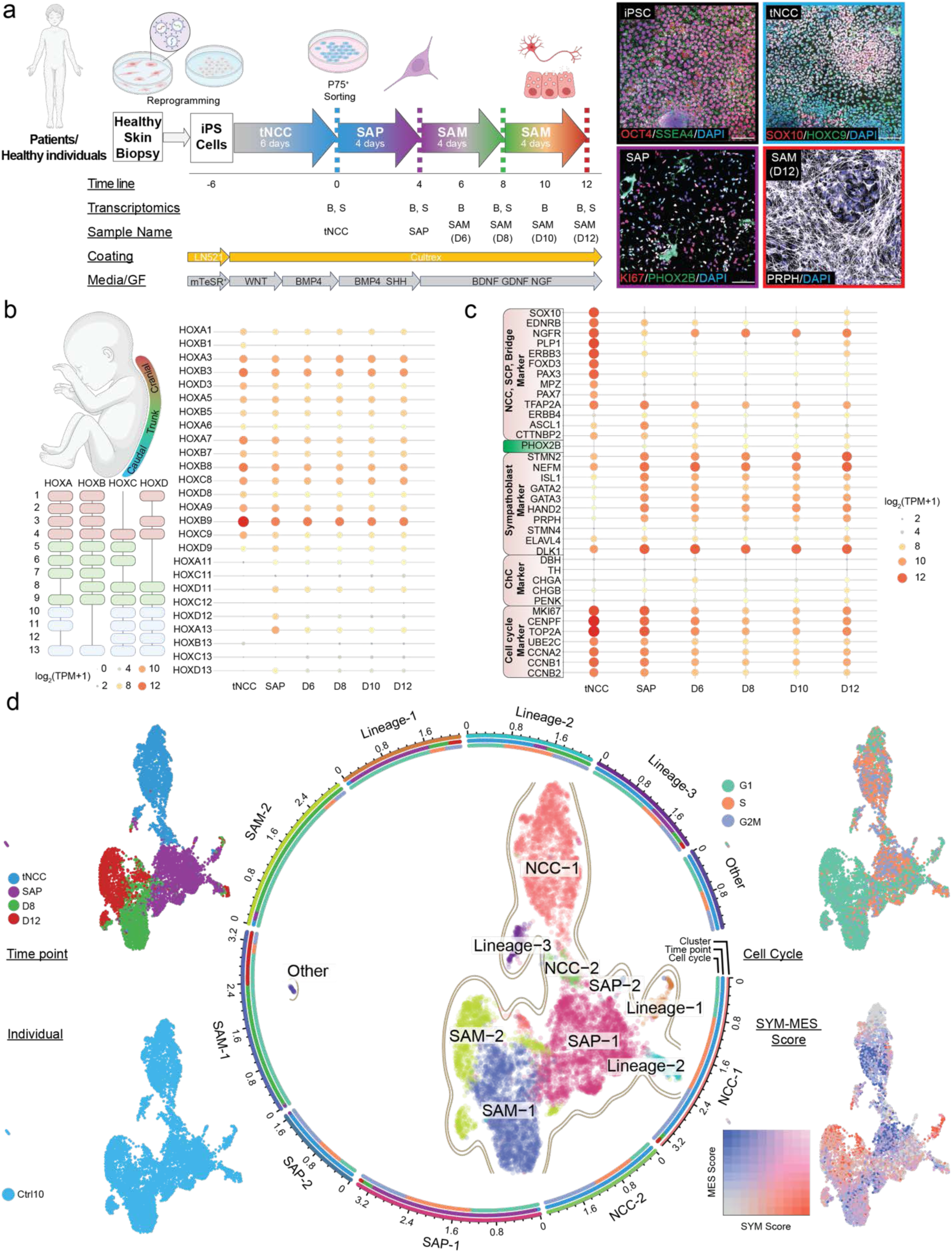
Stepwise differentiation of iPSCs into the sympathoadrenal lineage. **a,** Schematic representation of the differentiation protocol. The first row shows the transition from iPSC reprogramming to tNCC and subsequently to sympathoadrenal (SA) differentiation. Dashed lines of different colors indicate time points subjected to single-cell RNA sequencing, consistently applied throughout this study. The second row defines the project time line, with the tNCC stage designated as day 0. The third row lists bulk (B) or single-cell (S) RNA-seq performed at each collection point, and the fourth row indicates the names used for subsequent analyses. Rows five and six show the plate-coating substrates and growth factors used in culture. Representative merged immunofluorescence images (right) show iPSCs (OCT4, SSEA4), tNCCs (HOXC9, SOX10), sympathoadrenal progenitors (SAP; PHOX2B, KI67), and sympathoadrenal mature cells (SAM, day 12; PRPH). Nuclei were counterstained with DAPI. Scale bar, 100 µm. Corresponding single-channel images are available in the Supplementary Figures. **b,** Expression of selected HOX family genes was analyzed to assess anterior–posterior identity. Color-coded cranial (pink), trunk (green), and caudal markers (blue). The bubble plot illustrates HOX gene expression levels across collection time points (log₂[TPM+1]). **c,** Bubble plot showing expression of selected markers of NCC-to-SA lineage differentiation, based on published datasets of human adrenal medulla development. Values are shown as average log₂ (TPM+1). **d,** UMAP visualization of single-cell RNA-seq data from Ctrl cell differentiation. Cells were assigned to ten transcriptional clusters, indicated by colors and annotated (as detailed in the Methods section) by collection time point (upper left), individual identity (lower left), and cell-cycle phase (upper right), and SYM-MES score (lower right). The circular plot surrounding the UMAP shows cluster composition by time point and cell-cycle status, arranged clockwise by time point numbers indicate the log₁₀-scaled angular positions of ordered cell groups along the circular layout (middle).

To further resolve cell types in our model, we performed single-cell RNA-sequencing (scRNASeq) on Ctrl10 cells from three independent inductions, with four matched differentiation time points collected in each induction. (Fig. 1d; Extended Data Fig. 3–4). After quality control, 9,064 high-quality cells were retained for downstream analysis. These cells clustered into 10 transcriptionally distinct groups (Extended Data Fig. a-b). Cluster annotation was defined using multiple criteria, including developmental stage, cell-cycle state, marker gene expression, differential expressed gene (DEG) distinguishing NCC-derived mesenchymal (MES) and sympathoadrenal lineages (SYM) in mouse embryonic development^44^, pseudotime ordering, and cluster-specific transcriptional programs (Fig. 1d; Extended Data Fig. 3e–f, 4c–g, Supplementary Data 2). Clusters were annotated and defined by increased expression of markers genes as NCC-1 (primary NCCs) (*SOX10, PAX3*, and *ERBB3*), NCC-2 (high mitochondrial activity) (*MT-CO3*, *MT-ATP*, and *PAX3*), SAP-1 (early progenitors) (*HAND1*, *PTCH1*, and *ERBB3*), SAP-2 (*ATF3* and *GMFB*), SAM-1 (mesenchymal-like) (*FN1* and *COL1A1*), SAM-2 (sympathetic neuron-like) (*PLK2*, *SLIT3*, and *SYNPO*), Lineage-1 (mature SA lineage) (*PHOX2B*, *PHOX2A*, and *ASCL1*), Lineage-2 (immature SA lineage) (*MOXD1* and *POSTN*), Lineage-3 (neuroectodermal lineage)( *SOX2*, *MAP2*, and *GAP43*), and Other (*CD34*, *CDH5*, and *VIM*) (Fig. 1d, Supplementary Data 3).

Integration with published scRNA-seq data from hESC–derived SA models^26^ demonstrated strong similarity across corresponding developmental trajectories (Extended Data Fig 4e). Correlation analysis showed that our tNCCs most closely resembled intermediate-stage cells (D6-D9 NCC stage in the reference dataset), while later SA populations aligned well with terminally differentiated, more mature cells (Extended Data Fig. 4e–f).

Together, these results demonstrate that our model reliably captures key features of human SA lineage development, reflected both in *in vivo* developmental features and in strong alignment with existing *in vitro* models.

### Transcriptomic Profiling Reveals Divergent Developmental Trajectories in Patient vs. Ctrl Cells

Next we generated ALK R1275Q mutant iPSC lines by reprogramming healthy skin fibroblasts from three NB patients carrying germline ALK R1275Q mutations (NB1, NB2, NB4) (Fig. 1a, Fig. 2a, Extended Data Fig. 1a and c)^45^, hereafter these iPSC lines and their differentiated derivatives collectively referred as *patient-derived cells*. The patient iPSCs passed pluripotency quality control assays and retained the NM_004304.5 (*ALK*): c.3824G>A substitution in exon 25, corresponding to the ALK R1275Q mutation (Extended Data Fig. 1b and 1d). This mutation, located within the tyrosine kinase domain of ALK, induces a conformational change which is known to alter receptor activation (Extended Data Fig. 1c)^10,11,46^.

**Figure 2.**
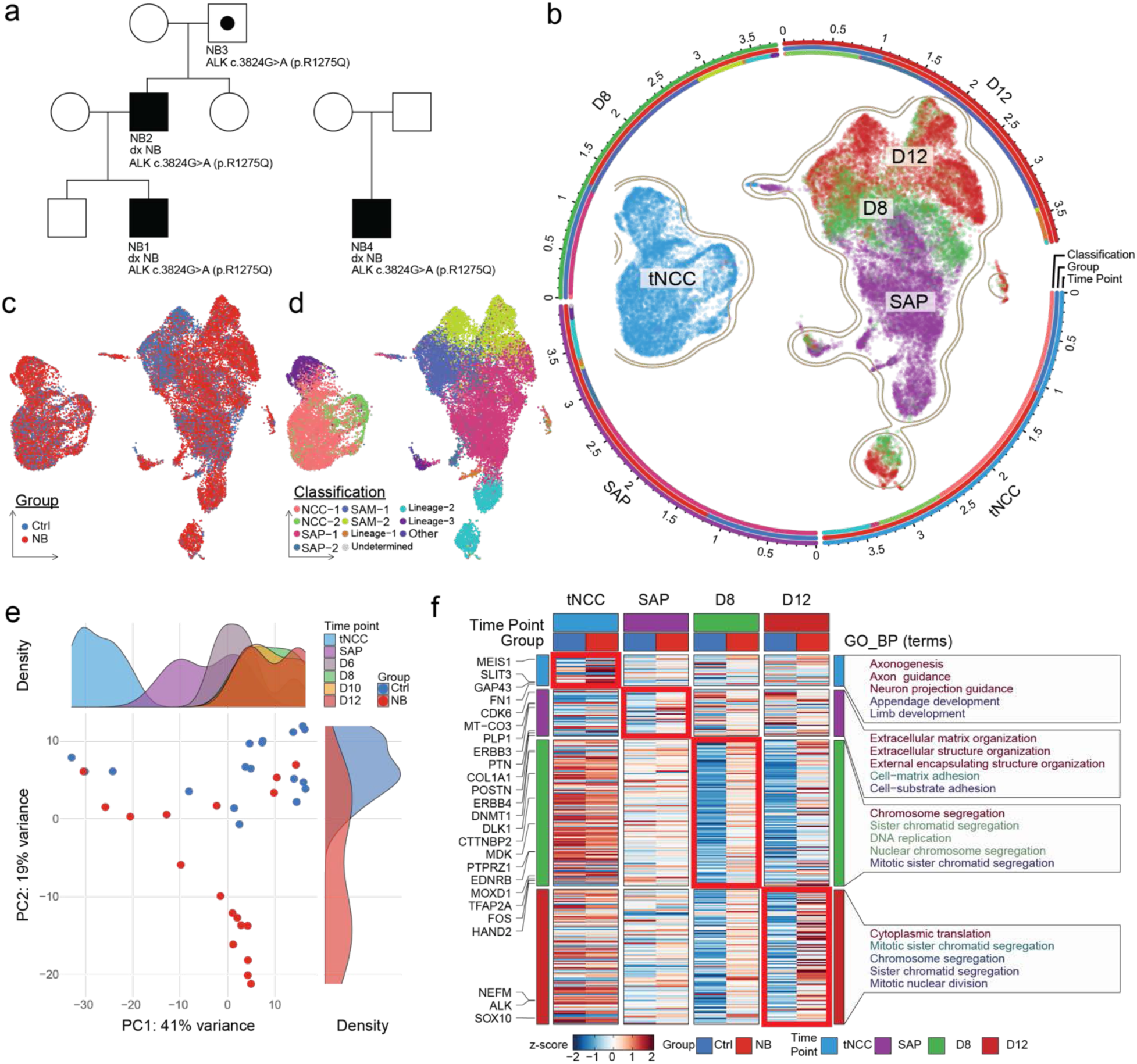
Single-cell transcriptomes distinguish patient-derived from Ctrl lines. **a,** Pedigree of patients included in this study. The familial NB cases NB1, NB2, and NB3 carry a germline ALK R1275Q mutation. NB1 and NB2 were diagnosed with NB, while NB3 never developed NB. NB4 represents a NB case with a *de novo* germline ALK R1275Q mutation. Patient-derived fibroblasts were reprogrammed into iPSCs: NB1, NB2, and NB4 were successfully reprogrammed, whereas NB3 fibroblasts failed to reprogram due to limited proliferative capacity. **b-d,** UMAP visualization of single-cell RNA-seq data from Ctrl and patient-derived lines, colored by collection time point. Circular plots surrounding the UMAP indicate cell composition by group (Ctrl versus NB), individual identity (Ctrl10, NB1, NB2, NB4), cell-cycle status, and classification. Classification was performed by mapping all cells onto Ctrl-derived reference clusters using k-nearest neighbor (KNN) prediction. Cells identities colored by group (Ctrl versus NB) and by predicted classification based on Ctrl-derived reference clusters (KNN prediction), are also displayed in the inserts c and d, respectively. **e,** Principal component analysis (PCA) of bulk RNA-seq data across time points. Density histograms display the distribution of samples by collection time point (top) and by group (Ctrl or NB, right), using kernel density estimation (KDE) for smooth approximation of sample distributions. **f,** Heatmap of z-scored mean expression values of differentially expressed genes (DEGs) upregulated in NB compared to Ctrl lines across time points in single-cell RNA-seq data. The color scale represents high (red) to low (blue) expression. Red bars mark the time points used for DEG calculation. Genes associated with SA lineage differentiation are labeled on the left. Gene ontology biological process (GO_BP) enrichment analysis of DEGs is shown on the right, with the top five terms (if applicable) displayed.

Building on our validated SA lineage model system, patient iPSCs were differentiated, and we performed bulk RNA-seq and scRNA-seq and integrated with the Ctrl dataset, using the previously obtained Ctrl clustering as a reference to compare differentiation trajectories (Fig. 2b, Supplementary Fig. 6). Samples were multiplexed at a 1:1:1:1 ratio across groups, resulting in an overall 3:1 distribution between patient-derived and Ctrl cells, and 1:1:1:1 at the individual sample level (Supplementary Fig. 6a-b). Integration of the scRNA-seq data yielded 30,654 high-quality cells. While integration preserved the distribution of differentiation time points (NCC, SAP, D8, D12) (Fig. 2b), it also revealed distinct differences between Ctrl and patient-derived cells (Fig. 2c, Extended Data Fig. 5b and d, Supplementary Figure 6a-b). Clustering analysis identified 14 transcriptionally distinct clusters (C0–C13) (Extended Data Fig. 5a, Supplementary Fig. 6c-d, Supplementary Data 4). Clusters comprised cells from multiple time points and sample origins, with varying relative proportions across clusters (Extended Data Fig. 5b–e).

Following integration of the control and iPSC-derived samples, we revisited the collection time points represented in each cluster (Fig. 2d). Cells collected at the NCC stage belonged primarily to the clusters NCC-1 (C2) and NCC-2 (C8), with some cells following the Lineage-3 trajectory and fewer cells from both Ctrl and patients following SAP and later-stage clusters (C3, C10). Most cells collected at the SAP time point belonged to the SAP-1 clusters (C0 and C6), while a subset of patient-derived SAP cells (C7) mapped to Lineage-2, representing immature SA populations. Ctrl cells collected at D8 and D12, were primarily grouped in SAM-1 and SAM-2 (C3), whereas patient-derived cells formed three distinct clusters (C4, C5, C9), resembling SAP-1, SAM-2 (sympathetic neuron-like), and immature SA Lineage-2, respectively. Mature SA lineage cells were represented by cluster C11 in the integrated dataset (Fig. 2d; Extended Data Fig. 5a). The distribution of cell cycle stages in the embedding suggests differences in proliferative capacity among cells during differentiation (Extended Data Fig. 5c). Notably, the divergence between control and patient-derived cells became most apparent after commitment to the SA stage. Supporting this observation, principal component analysis (PCA) of bulk RNA-sequenced samples revealed minor transcriptional differences between Ctrl and patient-derived cells at the NCC stage, that progressively increased through SAP and SAM stages (Fig. 2e). Single cell DEGs and enrichment analyses confirmed that the most pronounced differences emerged after the SAP stage, especially at D8 and D12, where patient-derived cells showed enrichment in gene programs related to mitosis and proliferation (Figure 2f, Supplementary Data 5).

Overall, these results highlight substantial heterogeneity and divergent differentiation trajectories between patient-derived and Ctrl cells, particularly during the transition from SAP to SAM. Patient-derived cells followed a perturbed developmental path, forming distinct clusters that reflected both normal and aberrant SA lineage states.

### Patient-derived Cells Remain in a Highly Proliferative and Less Differentiated State

To explore the cause of this altered trajectory, we next asked whether differences in proliferation might contribute to the delayed differentiation in patient-derived cells. Cell cycle state analysis of the single cell data showed that both Ctrl and patient-derived cells where actively proliferating in NCC and SAP, with >50% in S and G2/M phase. By D8 and D12, however, only a small fraction of Ctrl cells remained cycling, whereas a significantly higher proportion of patient-derived cells persisted in S and G2/M (Fig. 3a; Extended Data Fig. 6a, Supplementary Data 1). Further expression analysis of the bulk RNAseq data confirmed a sustained expression of cell-cycle genes^47^ in patient-derived cells, that was significantly higher than that of Ctrl cells across SA stages (Fig. 3b; Extended Data Fig. 6a). From D8 onwards, gene set enrichment analysis (GSEA) further showed enrichment of proliferation- and protein-synthesis programs, including E2F, MYC, mTORC1, and G2/M checkpoint gene sets (Extended Data Fig. 5f). Consistently, monitoring cell confluency in culture revealed an increase in patient-derived cells compared with Ctrl cells at every stage, from SAP to SAM D12 (Extended Data Fig. 6b).

**Figure 3.**
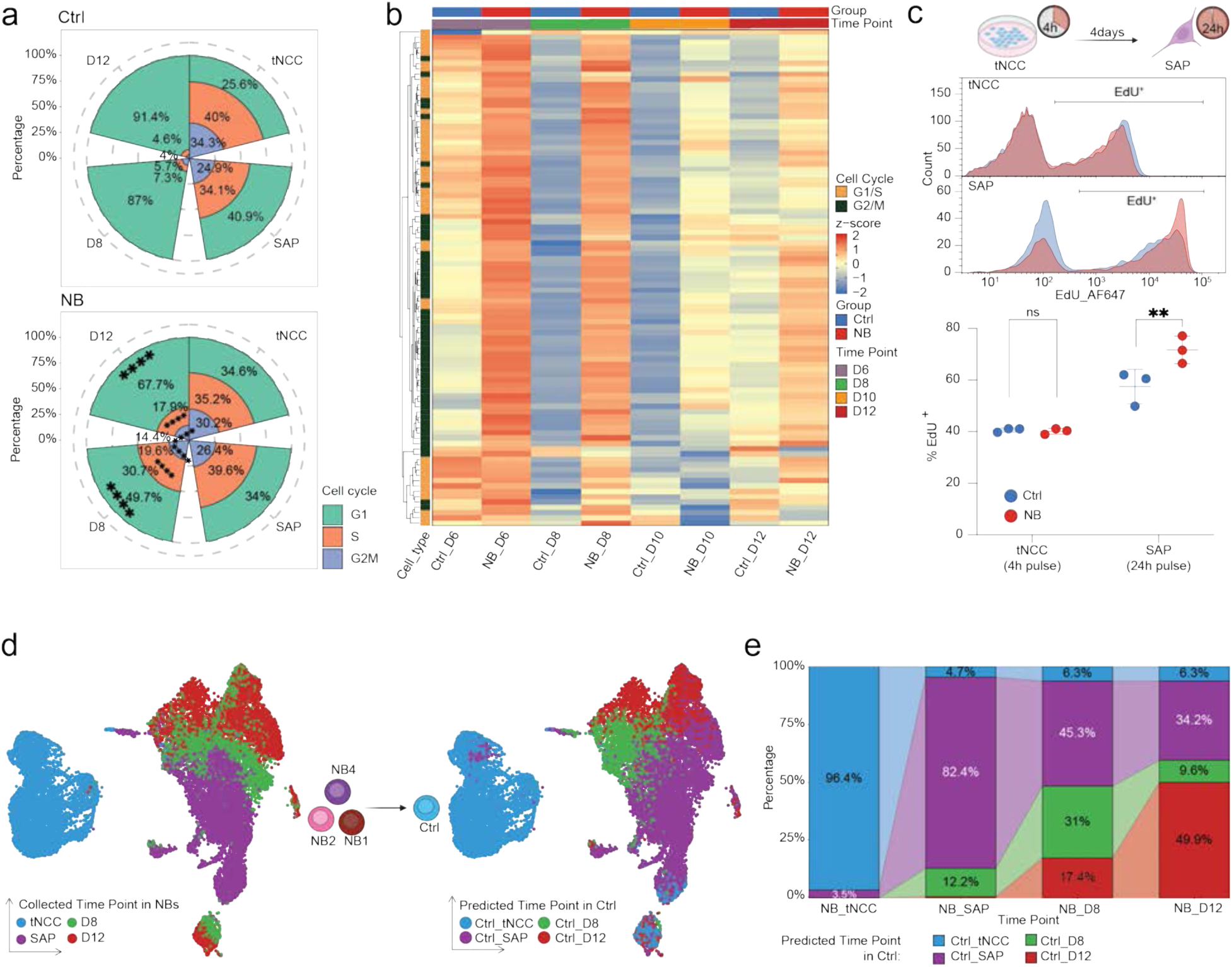
Patient-derived cells display sustained proliferation and impaired differentiation. **a,** Rose plots showing the proportion of cells in each cell-cycle phase (G1, S, G2/M) at four collection time points, based on single-cell RNA-seq data. Results are shown for Ctrl lines (top) and patient-derived lines (bottom). Comparisons were performed using Chi-square tests with FDR correction for multiple testing. Ctrl vs NB in G1, S, and G2M at D8 and D12, *FDR* < 0.0001 = ****. **b,** Heatmap of z-scored expression of cell-cycle genes^47^ across collection time points during SA lineage commitment. Group and time point annotations are shown on top of the plot. **c,** EdU incorporation assay demonstrating proliferation in Ctrl (Ctrl7, Ctrl 10 and Ctrl14 in blue) and patient-derived (NB1, NB2 and NB4 in red) lines at the NCC and SAP stages. Schematic (top) illustrates a 4h EdU pulse in tNCCs and a 24h pulse in SAP. Representative histograms of EdU-positive cells measured by flow cytometry (middle) and quantification of EdU+ cell percentages with statistical comparisons (bottom) are shown. Quantification of EdU⁺ cell percentages is shown as mean ± s.d. from independent biological replicates. Statistical significance was determined using a two-tailed Student’s *t*-test. Ctrl vs NB, in SAP, *P* = 0.0075 = **. **d,** UMAP visualization of patient-derived cells only (left), colored by collection time point, mapped onto Ctrl reference time points using KNN prediction. **e,** Bar plot showing the proportion of patient-derived cells assigned to each Ctrl reference time point.

Consistent with these findings, 5-ethynyl-2’-deoxyuridine (EdU) incorporation assays were performed at both the NCC and SAP stages. All three Ctrl and three patient-derived lines showed similar proliferation at the NCC stage. In contrast, patient-derived cells exhibited a significantly higher proportion of EdU+ cells than Ctrl cells at the SAP stage (Fig. 3c), confirming increased proliferation.

To compare differentiation outcomes between collected- and integration-predicted time points, patient-derived cells were extracted from the integrated single-cell dataset and mapped to the original Ctrl cells clustering as a reference. On the resulting embeddings, patient-derived and Ctrl cells showed no notable differences at the NCC stage after prediction. However, after prediction, patient-derived cells displayed a marked expansion of the SAP stage. Notably, a cluster composed mainly of patient-derived cells from D8 and D12 was mapped to the NCC stage in Ctrl cells (Fig. 3d), indicating a delayed differentiation.

Analysis of predicted cell proportions supported this observation. More than 96% of patient-derived cells collected at NCCs stage mapped to NCC time points in the Ctrl reference, and >80% of patient SAP cells did to SAP stage. However, over half of patient-derived cells remained in NCC or SAP states at D8, and by D12, more than half still resembled NCC, SAP, or D8 stages (Fig. 3e).

Finally, label transfer analysis from patient-derived cells to Ctrl cells confirmed these findings (as detailed in methods). While patient NCC and SAP cells were largely assigned to corresponding Ctrl stages, a subset of patient D8 and D12 cells was reassigned to earlier NCC or SAP stages of Ctrl cells (Extended Data Fig. 7), further supporting a delay in differentiation.

In summary, patient-derived cells remain more proliferative and less differentiated than Ctrl cells after commitment to the SA lineage, resulting in an accumulation of immature SA-like populations at later stages.

### ALK R1275Q Mutation Leads to Elevated *ALK* Expression and Increased Downstream Signaling Activity

Previous studies have reported that *ALK* mutations are associated with elevated *ALK* mRNA and protein expression in patient cohorts ^10^. In, particular, R1275Q results in gain-of-function kinase activity^14^. Given that patient-derived cells remained highly proliferative and less differentiated after SA commitment, we next asked whether this phenotype might be linked to altered *ALK* expression and signaling activity. To test whether the R1275Q mutation has similar effects in our system, we analyzed *ALK* expression dynamics across differentiation stages.

ALK expression was detectable at the NCC stage and remained relatively stable through the SAP stage. In Ctrl lines, ALK expression gradually declined during SAP-to-SAM differentiation. In contrast, patient-derived lines showed increasing ALK expression over time, peaking at later SA stages (Fig. 4a–b). Using Reactome pathway signatures (Reactome: R-HSA-201556, Supplementary Data 1), we calculated an ALK activity score at the single-cell level. From the SAP stage onward, patient-derived cells exhibited both higher ALK expression and elevated ALK activity compared with Ctrl cells (Fig. 4b).

**Figure 4.**
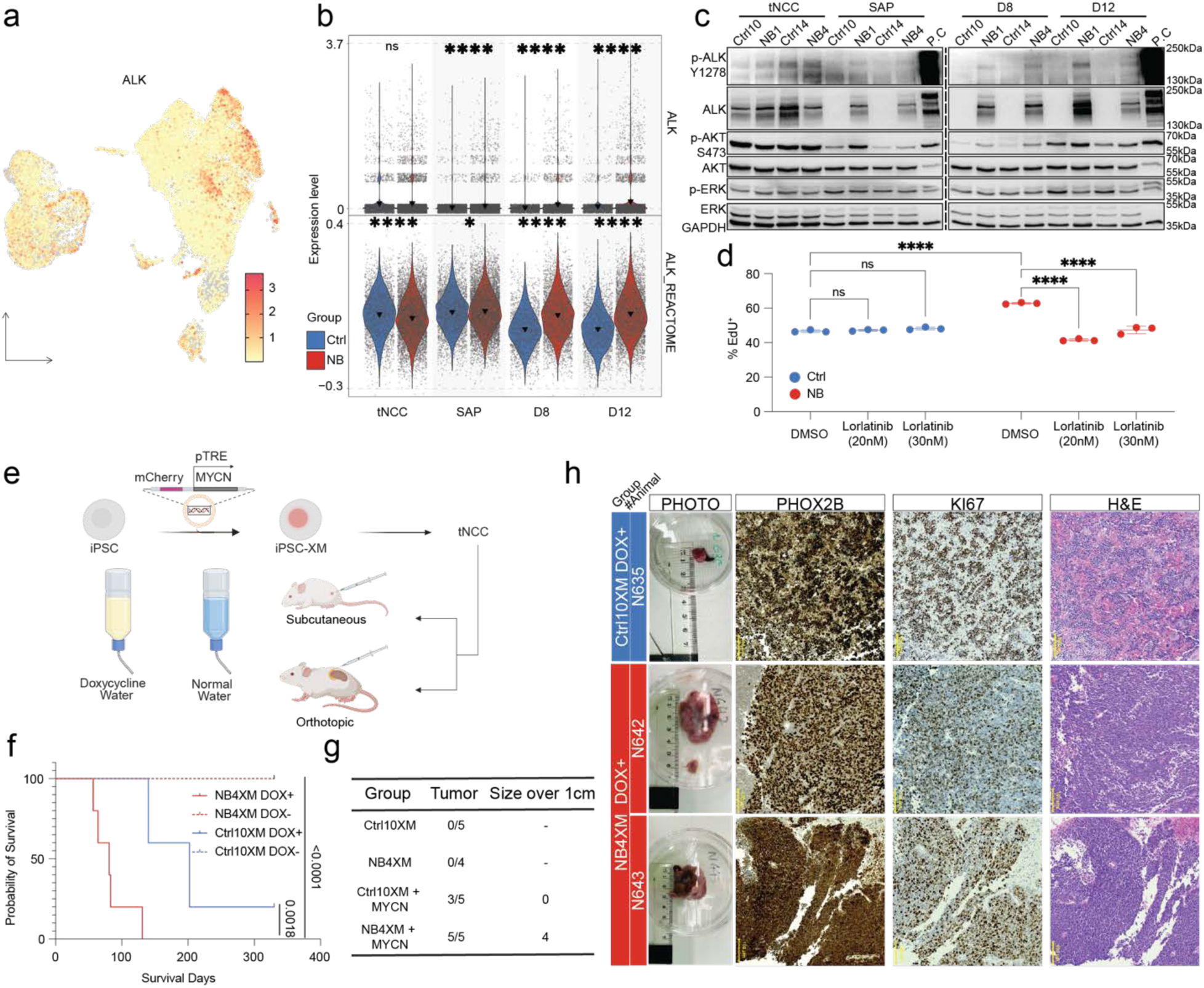
The ALK R1275Q mutation enhances downstream signaling but does not drive tumorigenesis alone. **a,** UMAP of all samples (as in Fig. 2b), colored by ALK expression. **b,** Violin plots of ALK expression (top) and ALK activity scores based on Reactome pathways (bottom) across time points in Ctrl and patient lines (computed as detailed in Methods). Comparisons were performed using the Wilcoxon test. ALK expression, Ctrl vs NB in SAP, D8, and D12, *P* < 0.0001 = ****; ALK Reactome score, Ctrl and NB in tNCC, D8, and D12, *P <* 0.0001 = ****, in SAP, *P =* 0.036 = *. Black triangles indicate mean values. **c,** Immunoblot analysis of protein lysates collected at NCC, SAP, day 8 (D8), and day 12 (D12) from two Ctrl lines and two patient-derived lines. Total and phosphorylated ALK, AKT, and ERK1/2 were probed. GAPDH was used as a loading control. P.C., positive control: NB-1 neuroblastoma cell line (CVCL_1440) treated with ALKAL2^19^. **d,** Quantification of EdU+ cells at the SAP stage after Lorlatinib treatment of both Ctrl (Ctrl 7, Ctrl10, and Ctrl14 in blue) and patient-derived (NB1, NB2, and NB4 in red) lines. Quantification of EdU⁺ cell percentages is shown as mean ± s.d. from independent biological replicates. Statistical significance was assessed by one-way ANOVA followed by Šídák’s multiple comparisons test. In DMSO, Ctrl vs NB, *P <* 0.0001 = ****; in NB, lorlatinib (20nM) vs DMSO, *P <* 0.0001 = ****; Lorlatinib (30nM) vs DMSO, *P <* 0.0001 = ****. **e,** Schematic of *in vivo* experimental design. iPSCs were stably transfected with a vector encoding constitutively expressed nuclear mCherry and doxycycline-inducible *MYCN*. Differentiated tNCCs were injected subcutaneously or orthotopically into the adrenal gland. MYCN was activated by mice receiving doxycycline (200 μg/ml) in drinking water. **f,** Kaplan–Meier survival curves from the subcutaneous injection model. Solid line, mice treated with doxycycline; dashed line, untreated controls. Significance was assessed by Cox proportional hazards test. **g,** Table summarizing results of the orthotopic injection experiment. Tumor size is reported as the longest axis of the tumor. **h,** Histological analysis of tumors from orthotopically injected mice. Gross tumor morphology is shown (with ruler indicating size, top). Representative sections were stained for PHOX2B, KI67, and hematoxylin and eosin (H&E) in the same tumor region. Scale bar, 100 µm.

In agreement with these transcriptional changes, ALK and phosphorylated ALK (pALK) protein levels were similar in control and patient-derived cells at the NCC stage. Interestingly, on SA differentiation, ALK and pALK levels rapidly declined in control cells but remained high in patient-derived cells. In addition, phosphorylated AKT levels increased in patient-derived cells at the SAP and D8 stages, suggesting enhanced ALK downstream signaling activity^48^. In contrast, both total and phosphorylated ERK1/2 levels were similar between control and patient-derived lines across differentiation (Fig. 4c).

To test whether the enhanced proliferation observed in patient-derived cells was ALK-dependent, we assessed EdU incorporation at the SAP stage following treatment with lorlatinib, a third-generation ALK inhibitor. Lorlatinib treatment reduced the fraction of EdU+ patient-derived cells at the SAP stage from 62.8 ± 0.5% to 41.5 ± 0.67% (20nM) or 47.4 ± 2.15% (30nM), approximating the baseline level of Ctrl cells. In contrast, Ctrl cells remained largely unaffected (47.7 ± 0.77% EdU+ with or 46.7 ± 0.69% without treatment), indicating that the mutation-driven proliferative component is ALK-dependent, while a baseline proliferative fraction persists independently of ALK activity. Increasing lorlatinib concentrations did not further reduce EdU incorporation, suggesting that only the ALK-dependent proliferative subset is drug-sensitive (Fig. 4d). Thus, only the increased proliferation in patient-derived cells is ALK-mutation-dependent, while baseline proliferation remains unchanged. Together, these findings demonstrate that the ALK R1275Q mutation sustains ALK expression and signaling activity during SA differentiation, driving aberrant proliferation that can be effectively suppressed by pharmacological ALK inhibition.

### ALK Mutation Creates a Permissive but Non-Tumorigenic State for Oncogenic Events

To assess if the increase in proliferation from the ALK R1275Q mutation alone drives tumorigenesis, we transplanted Ctrl and patient tNCC into immunocompromised mice, both subcutaneously and orthotopically (Fig. 4e). No tumors formed under these conditions (Fig. 4f–g), consistent with previous findings suggesting that ALK mutations require cooperating oncogenic drivers like MYCN amplification^20,49^. To test this, we engineered two Ctrl (Ctrl10XM, Ctrl14XM) and two patient iPSC lines (NB1XM, NB4XM) with a doxycycline-inducible MYCN construct (Extended Data Fig. 8a-b). Interestingly, MYCN activation altered differentiation dynamics *in vitro*, with a stronger effect in patient-derived cells (Extended Data Fig. 8c). Consistently, upon MYCN induction, mice injected with patient-derived tNCC developed larger and faster-growing tumors compared to Ctrl (Fig. 4e-g; Extended Data Fig. 8d). Histopathological analysis confirmed neuroblastoma-like features, including “small blue round cell” morphology and expression of KI67 and PHOX2B, consistent with proliferative neuroblastoma cells of sympathoadrenal origin.

These in vivo experiments demonstrate that while ALK R1275Q alone is not tumorigenic, it creates a permissive state that facilitates oncogenic cooperation and accelerates tumor initiation *in vivo*.

### Patient-derived Cells Retain a Proliferative SCP-like Population within the SA Lineage

Given that the increase in proliferation in patient-derived cells appeared upon commitment to the SA lineage, and that ALK inhibition suppressed this increase but not basal proliferation, we asked whether a specific subpopulation within the SAP stage is susceptible to mutant ALK activity. To investigate this, we performed subclustering of SAP-stage cells (Fig. 5a), identifying twelve subclusters (SAP_0–SAP_11) (Fig. 5b; Extended Data Fig. 9a–d). Among these, SAP_5 stood out for its enrichment in patient-derived cells, spanning all cell-cycle phases, and elevated proliferative activity (Fig. 5c–d). Importantly, SAP_5 also showed the highest ALK activity scores, with patient-derived cells consistently exhibiting stronger ALK activity than Ctrl cells (Fig. 5e–f).

**Figure 5.**
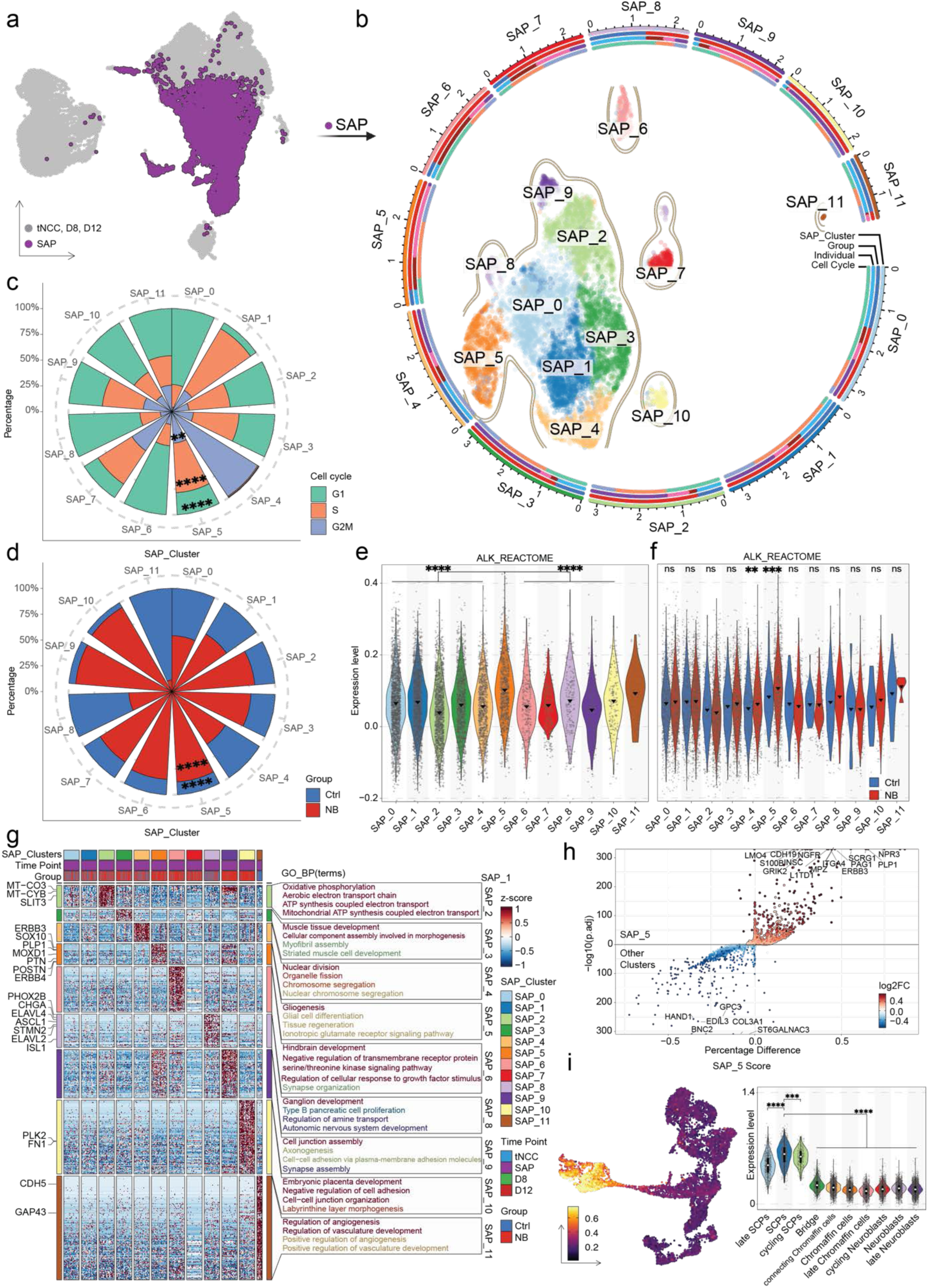
Identification of a proliferative SCP-like subpopulation in patient-derived cells. **a,** UMAP of all samples (as in Fig. 2b), highlighting SAP-stage cells selected for subclustering. **b,** UMAP of SAP-stage cells following subcluster analysis. Twelve clusters (SAP_0-SAP_11) were identified and are shown in distinct colors. Circular plots surrounding the UMAP indicate cell composition by group (Ctrl versus NB), individual identity (Ctrl10, NB1, NB2, NB4), and cell-cycle status (computed as detailed in Methods). The color used in the circle is the same as in Extended Data Fig.9a-c. **c-d,** Rose plots showing the proportion of cells in each cell-cycle phase across SAP subclusters (c) and the proportion of Ctrl versus patient line cells within each SAP subcluster (d). Comparisons were performed using Chi-square tests with FDR correction for multiple testing. G2M, SAP_5 vs other, *FDR =* 0.00109 =**, G1 and S, SAP_5 vs other, *FDR* < 0.0001 = ****. Ctrl and NB, SAP_5 vs other, *FDR <* 0.0001 = ****. **e-f,** Violin plots showing the distribution of ALK activity scores in each SAP subclusters (e) and the distribution of ALK activity scores in Ctrl versus patient lines within each SAP subclusters (f). Black triangles indicate mean values. Statistical significance was assessed using the Wilcoxon test. ALK Reactome score between SAP_5 and other groups, *P* < 0.0001 = ****. ALK Reactome score, Ctrl vs NB in SAP_4, *P* < 0.088 = **; in SAP_5, *P* = 0.00054 = ***. **g,** Heatmap of z-scored expression values of DEGs upregulated in each SAP subcluster. Color scale represents high (red) to low (blue) expression. Genes associated with SA lineage differentiation are labeled on the left. Enrichment analysis of DEGs in GO-defined Biological Processes is shown on the right, with the top five terms (if applicable) displayed. Color denotes FDR values (red to blue, as detailed in methods). **h,** Scatter plot showing DEGs between the SAP_5 cluster and all other SAP clusters. The y-axis shows –log₁₀(adjusted P value), and the x-axis shows the percentage difference between SAP_5 and other clusters. Points are colored by log₂ fold change, with genes upregulated in SAP_5 shown in red and downregulated in blue. **i,** UMAP of published human adrenal medulla single-cell data^42^ colored by SAP_5 module score (yellow, high; purple, low). Violin plots show the distribution of SAP_5 scores across clusters in this dataset. Statistical significance was assessed using the Wilcoxon test. SAP_5 score, SCP vs cycling SCP, *P* = 0.00016 = ***; SCP vs other groups, *P* < 0.0001 = ****.

Marker analysis revealed that SAP_5 expressed both SA lineage markers (*DPYSL2, RORB, GATA3, SOX4, MOXD1*, *POSTN*) and classical SCP-associated genes (*ERBB3*, *NGFR*, *PLP1*, *S100B, SOX10*), along with enrichment of gliogenesis-related terms (Fig. 5g–h, Supplementary Data 6). For SAP_5 signatures analysis, SAP_5 markers were defined as genes significantly upregulated relative to other SAP subclusters (adj. *P* < 0.05, log₂FC > 0.5). Consistently, SAP_5 signature (indicating high-relative expression of upregulated SAP_5 markers) was significantly higher in SCPs across two independent human embryonic adrenal medulla datasets (Fig. 5i; Extended Data Fig. 9e–h)^42,43^. SCPs have been identified as an NCC-derived progenitor population that can contribute to sympathoadrenal cells^50^. We therefore assessed whether the SAP_5 transcriptional signature is more characteristic of an SCP-like or an early NCC-like state. To clarify this, we scored Ctrl-derived tNCC and SA lineage populations, originally built as a reference and included in Figure 1d. SAP_5 signature was significantly higher in cells of Lineage-2 (immature SA) compared with cells in tNCC clusters (NCC-1, NCC-2, Fig. 6a). In support of these results, a cross-projection analysis (as detailed in Methods) showed that both Ctrl- and patient-derived SAP_5 cells mapped to the Lineage-2 cluster of the original Ctrl-embedding reference (shown in Figure 1d), with patient-derived cells being overrepresented (Fig. 6b–c). Importantly, this suggests that SAP_5 is not a transient or residual neural crest-like cluster, but rather a distinct population that emerges during early sympathoadrenal differentiation. It is defined by the co-expression of SCP and SA markers and increases and persists longer in patient-derived cells.

**Figure 6.**
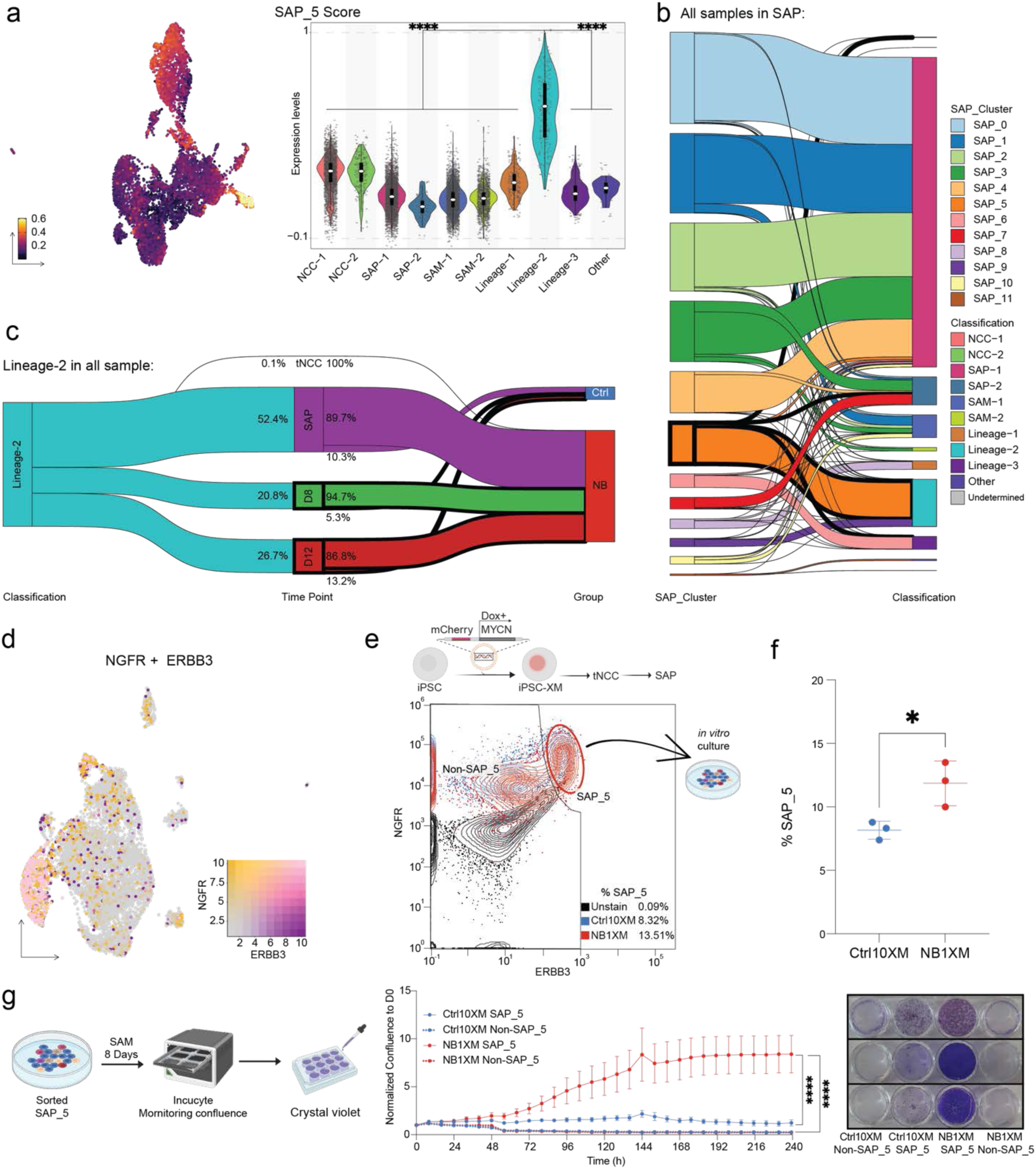
SAP_5 cells show increased transformation potential in patient-derived lines. **a,** UMAP of Ctrl cells only (as in Fig. 1d, middle), colored by SAP_5 module score (left, computed as detailed in Methods). Violin plots show the distribution of SAP_5 scores across clusters in the Ctrl dataset. Statistical significance was assessed through pairwise comparisons between Lineage-2 and each other groups of cells using the Wilcoxon test. SAP_5 score, Lineage-2 vs other groups, *P* < 0.0001 = ****. **b,** Sankey plot showing the transitions among SAP subclusters and KNN-predicted classifications at the SAP stage. The SAP_5 transition flow is highlighted with bold lines. **c,** Sankey plot showing the lineage transitions of Lineage-2 cells across time points and subsequently to Ctrl or patient lines. Numbers indicate the percentage within the preceding category (In each individual, Ctrl10 10%, NB1 24.8%, NB2 64.9%, and NB4 0.2%). **d,** UMAP of SAP-stage cells (as in Fig. 5b), colored by *NGFR* and *ERBB3* expression. NGFR-high cells are shown in yellow, ERBB3-high in purple, and double-positive cells in pink. **e,** Schematic of the SAP_5 enrichment strategy. iPSCs transfected with mCherry and doxycycline-inducible *MYCN* (as in Fig. 4e) were differentiated to the SAP stage via tNCCs, then stained for NGFR and ERBB3 for flow cytometry-based cell sorting. Representative contour plots (middle) show double-positive populations. Black indicates unstained negative control, blue indicates Ctrl cells, and red indicates patient-derived cells. The red circle highlights the SAP_5 population sorted for further culture. **f,** Quantification of sorted SAP_5 cells. Quantification of SAP_5 cell percentages is shown as mean ± s.d. from 3 independent inductions. Statistical significance was determined using a two-tailed Student’s *t*-test. SAP_5%, Ctrl vs NB, *P* = 0.0408 = *. **g,** Experimental workflow for SAP_5 cells. Sorted SAP_5 cells were cultured for 8 days in SAM medium, with or without *MYCN* induction, following the SAP-to-SAM protocol. Equal cell numbers were replated per well, and growth was monitored using IncuCyte. After 10 days, cells were collected for crystal violet staining (left). Growth curves (middle) show cells treated with doxycycline: solid lines represent SAP_5-sorted cells, dashed lines represent non-SAP_5 cells. Blue, Ctrl; red, patient-derived. Data are shown as mean ± s.e.m. NB1XM with Dox vs others, *P* < 0.0001 = ****. Representative crystal violet staining results from three independent experiments are shown (right).

In summary, SAP subclustering identified SAP_5 as a proliferative population enriched in patient-derived cells, transcriptionally resembling SCP-like cells. This population is characterized by high ALK activity and is persistence throughout SA differentiation. SAP_5 may represent a patient-enriched reservoir of proliferative, immature SA-like cells.

### SAP_5 Cells Exhibit Increased Susceptibility to Transformation, Especially in Patient-derived Cells

Having identified SAP_5 as a proliferative, ALK activity-high, SCP-like subpopulation enriched in patient-derived cells, we next investigated whether these cells are more susceptible to oncogenic transformation. To isolate SAP_5 cells, we analyzed cell surface marker combinations and identified the genes *ERBB3* and *NGFR* as highly expressed in this population (Fig. 6d, Extended Data Fig. 9i, Supplementary Data 7). Using these markers, we used flow cytometry to isolate SAP_5 and non-SAP_5 populations from Ctrl10 and NB1 (Fig. 6d–e, Extended Data Fig 9j, Supplementary Fig 7), confirmed by SAP_5 marker gene expression using RT-qPCR (Extended Data Fig. 9k). As predicted by transcriptomic data, SAP_5 cells were significantly more abundant in patient-derived SAP populations (Fig. 6f). Next, we cultured the sorted cells under differentiation conditions with doxycycline-induced MYCN-expression. SAP_5 cells showed significantly enhanced proliferative capacity compared to non-SAP_5 cells. This effect was observed in all SAP_5 cells but was significantly more pronounced in patient-derived SAP_5 cells (Fig. 6g). Thus, SAP_5 cells are likely to have increased susceptibility to clonogenic growth under oncogenic stress. Altogether, these findings suggest that patient-enriched SCP-like states may serve as a transformation-prone pool of cells during SA differentiation.

### SAP_5 Signatures Associate with ALK Activity, Age at Diagnosis, and Clinical Outcome in Patients

To extend our *in vitro* findings to patient data, we analyzed the Sequencing Quality Control neuroblastoma RNA-seq dataset (SEQC-NB, consisting of 498 NB patient tumors) using the SAP_5-derived transcriptional signature. Since ALK mutation status was not annotated, we first calculated ALK pathway activity scores (ALK_reactome score) as a functional proxy. Consistent with our iPSC-derived model, SAP_5 scores correlated positively with ALK activity across patient tumors (Fig. 7a). When stratified by MYCN status, MYCN-amplified tumors showed uniformly low SAP_5 scores, whereas non-MYCN–amplified tumors displayed significantly higher scores (Fig. 7b). K-means clustering (as detailed in methods) separated patients into SAP_5-high and SAP_5-low groups, with all MYCN-amplified tumors falling into the SAP_5-low group (Fig. 7c , Extended Data Fig. 10a).

**Figure 7.**
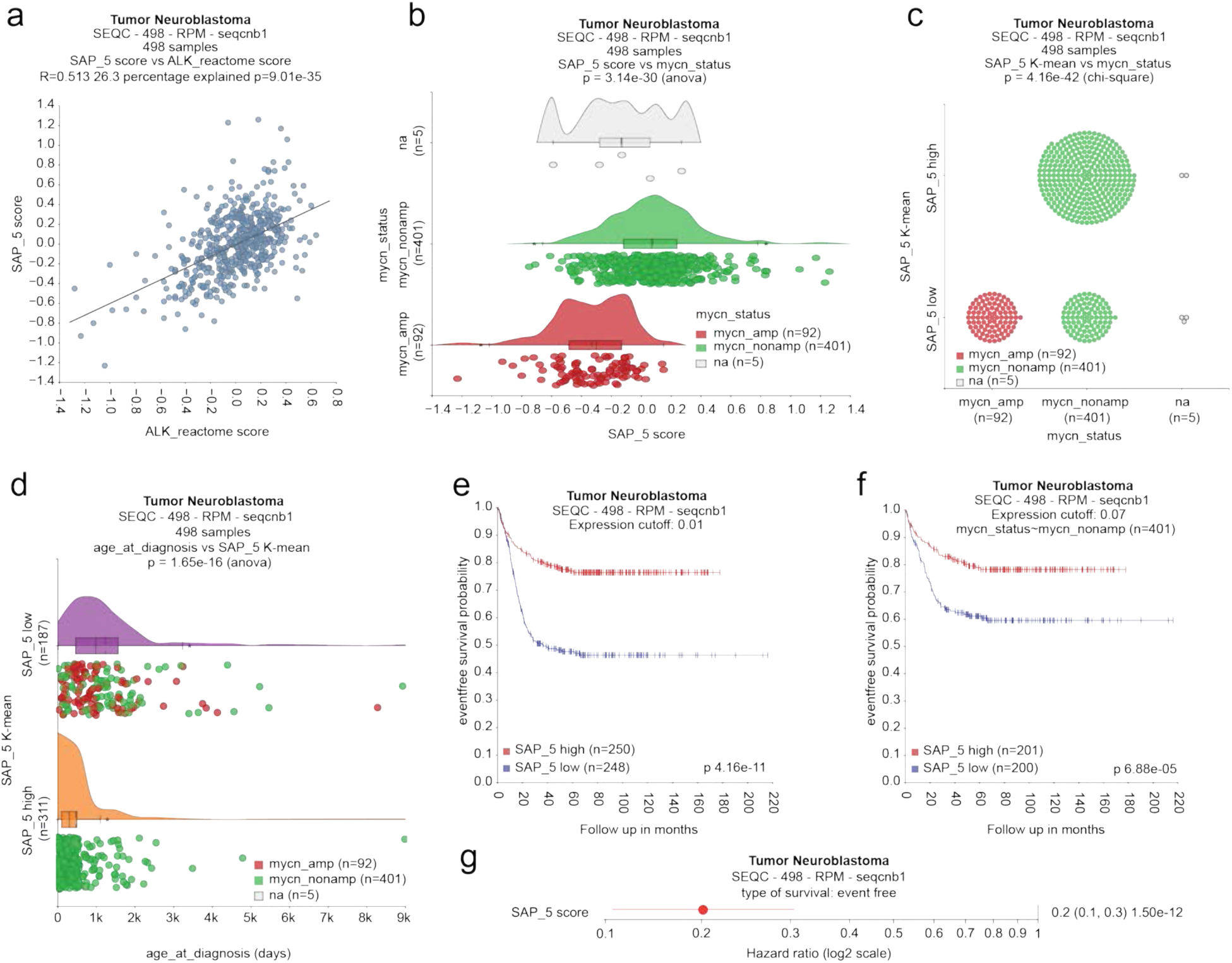
SAP_5 cells are associated with favorable prognosis and early diagnosis in the SEQC-NB patient cohort. **a,** Scatter plot showing the correlation between SAP_5 module scores and ALK Reactome scores in SEQC-NB data. Each dot represents a patient. The black line indicates the linear regression fit (R = 0.513). **b,** Raincloud plot showing SAP_5 scores stratified by MYCN status in 498 patients: MYCN-amplified (red, n = 92), MYCN non-amplified (green, n = 401), and unknown status (grey, n = 5). **c,** Box plot showing the distribution of MYCN status across SAP_5 subgroups defined by k-means clustering in SEQC-NB data. Colors as in b. **d,** Raincloud plot showing age at diagnosis (days) across SAP_5 subgroups defined by k-means clustering in SEQC-NB data. Colors as in b. **e–f,** Kaplan–Meier plots of event-free survival probability in SEQC-NB patients, stratified by SAP_5 scores (median split: high, red; low, blue). Analyses were performed in all patients (e) or restricted to MYCN non-amplified patients (f). **g,** Hazard ratios for SAP_5 scores in the SEQC-NB cohort with respect to event-free survival. Orange dots represent hazard ratios; horizontal lines indicate 95% confidence intervals (CIs). SAP_5 score had a hazard ratio of 0.20, where a hazard ratio < 1 indicates a reduced risk of adverse outcome (protective effect), and a hazard ratio > 1 indicates an increased risk (deleterious effect). (95% CI, 0.13–0.31).

We next examined clinical associations. Patients with high SAP_5 scores were diagnosed at younger ages compared with SAP_5-low patients (Fig. 7d), and survival analyses showed that higher SAP_5 scores predicted longer event-free and overall survival, independent of MYCN status (Fig. 7e–f; Extended Data Fig. 10b–e). Cox regression confirmed this effect (Fig. 7g).

In summary, SAP_5 signatures capture an ALK-associated SCP-like state that is enriched in younger, non-MYCN–amplified patients and correlates with a favorable prognosis. Oncogenic events such as MYCN amplification may rewire the transcriptome towards a malignant state, whereas in low-risk cases, the transcriptional identity remains closer to the original SAP_5 signature.

Together, our findings show that ALK R1275Q mutation sustains proliferative signaling and alters developmental trajectories through maintaining a SCP-like cell state. While the mutation alone is insufficient for tumorigenesis, it creates a permissive environment for other oncogenic hits. Analysis of NB patient tumor cohorts further supported the relevance of SAP_5, linking this SCP-like state to age at diagnosis and clinical outcome. This identifies a vulnerable window during sympathoadrenal differentiation that may underlie neuroblastoma initiation and inform future therapeutic strategies.

## Discussion

NB is an embryonic developmental cancer in which tumorigenesis has been linked to disrupted sympathoadrenal (SA) lineage differentiation^1,4^. By modeling this trajectory using human iPSC-derived SA lineage cells via trunk neural crest (tNCC) intermediates, we demonstrate that cells with ALK R1275Q mutation deviate from normal SA development. Specifically, these cells remain highly proliferative and incompletely differentiated following SA commitment, characterized by persistent cell-cycle activity and delayed acquisition of mature SA features. These findings are consistent with previous findings that aberrant proliferation and impaired lineage progression represent early hallmarks of neuroblastoma ^51,52^. However, our system uniquely provides direct insight into human developmental context, which has been largely inaccessible *in vivo*.

The failure of ALK-mutant cells to properly exit cell cycle suggests that the R1275Q mutation not only amplifies mitogenic signaling but also perturbs the balance between self-renewal and differentiation. Notably, the persistence of SCP-like cells shows that ALK mutations generate a developmental “stall point” where progenitor populations accumulate. This is consistent with murine and zebrafish studies showing that ALK mutations bias neural crest derivatives towards proliferation^53–56^. Our data extends these findings by demonstrating that this effect also occurs in human SA differentiation. Thus, the ALK R1275Q mutation functions less as a conventional oncogenic “switch” and more as a lineage-modifying factor that constrains cells in a proliferative, transformation-prone state. In line with this interpretation, our single-cell analysis revealed a persistent cluster (SAP_5) enriched in proliferative, immature SA-like cells, which was more pronounced in patient-derived cells than in Ctrl cells.

The question of NB cell-of-origin remains a matter of ongoing debate, with sympathoblasts, chromaffin cells, and SCPs all considered as candidate progenitor cell types^42,43,57–61^. Previous work has shown that copy number alterations that include 17q/1q gains, frequently detected in high-risk NB, can impair the specification of tNCC before SCP stage and enhance the tumorigenic effects of *MYCN*^26^. Furthermore, a *SOX2*⁺ progenitor-like population was previously reported in ALK-mutant iPSC models^34^, suggesting that ALK mutations can stabilize immature cell states. In our work, *SOX2* expression was also elevated within the SAP_5 subpopulation (Supplementary Data 6). However, SAP_5 was more distinctly defined by SCP-associated markers such as *ERBB3*, *NGFR*, *PLP1*, and *S100B*, together with enrichment of gliogenesis-related programs. In contrast to the previously described SOX2⁺ cells, SAP_5 populations were present in both control and patient-derived cells but were significantly more expanded and persistent in the latter. To us, this suggest that ALK mutations do not generate a *de novo* progenitor state but rather stabilize and amplify a normally transient progenitor pool. Functionally, SAP_5 cells had elevated ALK activity and enhanced susceptibility to MYCN-driven transformation, consistent with a transformation-prone progenitor state directly linking this subpopulation to proliferative imbalance and tumor initiation risk. Importantly, we do not claim that SAP_5 represents the definitive NB cell-of-origin; rather, our data support the view that ALK mutations act as an early “first hit,” extending the developmental window during which cells can acquire additional oncogenic events (“second hits”). The cell-of-origin may vary depending on the specific genetic lesion and the developmental context in which it arises. Together with previous studies, this supports a model in which NB arises not from a single uniform cell of origin, but from multiple progenitor states rendered permissive by distinct genetic lesions and developmental timing.

ALK functions as a receptor tyrosine kinase (RTK) that normally mediates proliferative signaling upon ligand binding. Mutations within its kinase domain, however, have been reported to drive constitutive downstream activation in a ligand-independent manner^10,11,46,62^. In our study, ALK-mutant cells maintained proliferative signaling beyond the neural crest stage, with patient-derived lines showing enhanced proliferative capacity compared with Ctrl lines. This extended proliferative state could be reduced to baseline by treatment with the ALK inhibitor Lorlatinib, underscoring the coexistence of both ALK-dependent and ALK-independent signaling during sympathoadrenal differentiation.

The cooperative role of ALK mutations with other proliferative stresses may help explain their contribution to NB initiation. Acting as an early “first hit,” ALK mutations extend the developmental time window during which additional oncogenic alterations (“second hits”) can be acquired. Cooperation between ALK mutations and MYCN has been demonstrated in GEMMs and hPSC-derived NCC xenografts^18,20,35,36^, typically using the potent ALK F1174L variant. In our study, the ALK R1275Q mutation behaved similarly: alone insufficient for tumor initiation, but together with MYCN markedly accelerating tumor formation. Previous studies have shown that mutant ALK can phosphorylate CHK1 at non-canonical sites that are not inhibited by ATR inhibitors, thereby perturbing p53 pathway activity^63^. Such mechanisms may help explain why ALK-mutant cells are more prone to transformation and are associated with more aggressive disease phenotypes compared with their wild-type counterparts. These mechanistic insights have direct implications for clinical manifestations and therapeutic responses.

From a clinical perspective, NB harboring only ALK mutations are relatively uncommon (∼8% with single somatic mutations and ∼10–14% when including amplifications)^10,11,15,46,64–67^. Such cases are usually accompanied by additional genetic alterations, including MYCN amplification, ATRX mutation, and recurrent chromosomal aberrations such as 2p gain, 17q gain, 11q loss, and 1p loss ^15,46^. Among the hotspot variants, ALK R1275Q is the most frequent, accounting for ∼45% of familial NB and ∼33% of sporadic cases ^46,68^. Unlike the F1174L mutation, which is strongly associated with MYCN amplified, high-risk NB, R1275Q is observed across both low- and high-risk groups. Moreover, across ages, the distribution of ALK mutation sites (including R1275Q) does not differ significantly^69^. These patterns suggest that ALK mutations act primarily to enhance the oncogenic activity of cooperating drivers, rather than serving as a sole initiator. Although only 1-2% of NB cases are familial, the estimated penetrance of germline ALK mutations approaches 50%^17^. This underscores the need for monitoring germline ALK carriers, where surveillance for secondary genetic “hits” may improve early detection and risk reduction.

The clinical correlations support our model that SAP_5 represents a transformation-prone progenitor state. In younger patients, this state likely reflects a differentiation defect driven by ALK activation, consistent with their better prognosis. In contrast, older patients frequently acquire additional oncogenic events, such as MYCN amplification, which cooperate with ALK-driven programs to promote more severe transformation and poor clinical outcome.

Therapeutically, ALK inhibitors such as the third-generation compound lorlatinib have improved responses in relapsed or refractory NB, with ∼30% objective response ratio as monotherapy and ∼60% in combination with chemotherapy^70^. Durable responses, sometimes exceeding three years, have been reported in ALK R1275Q patients^71^, underscoring their clinical value. Yet, limitations are evident. Our data point to heterogeneity within the SAP stage, which includes both control-and patient-derived progenitor-like cells with variable characteristics. While lorlatinib suppressed ALK-driven hyperproliferation, such intrinsic cellular diversity may contribute to variable or incomplete responses and, over time, to increased tumor heterogeneity. In particular, since ALK mutations can expand progenitor populations, these findings suggest that ALK inhibition alone may not be sufficient to fully control disease evolution. Rational combination strategies, for example, with ATR inhibitors to block compensatory signaling^63^, may therefore be necessary to achieve greater and more durable responses.

Looking forward, while SAP_5 represents a population of cells with transformation-prone features, the underlying regulatory networks, including transcriptional and epigenetic mechanisms, remain to be defined. Further studies will be essential to clarify how these programs are established and maintained. In addition, the iPSC-based model described here offers a platform for therapeutic exploration. This system could be leveraged for drug screening, as well as for evaluating innovative treatment strategies such as gene therapy or cell therapy. Together, these approaches may provide new avenues for developing effective interventions for neuroblastoma patients.

## Materials and Methods

### Ethics statement

All experiments involving human-derived material were conducted in accordance with ethical guidelines and approved by Swedish ethical review authority, EPN2009/1369/31/1, EPN 2012/208-31/3 with addendum 2012/856-32. Informed consent was obtained from all participating patients or their legal guardian. All animal procedures complied with Karolinska Institutet (KI) guidelines and were approved by the Stockholm North Ethical Committee for Animal Research (10025/23 with addendum 5503-2024).

### Animal experiment

Immunocompromised NSG mice (6–9 weeks; both sexes) were housed at Komparativ Medicin Annexet (KM-A), KI. Ctrl10XM or NB4XM tNCCs were transplanted either orthotopically into the left adrenal fat pad (2 × 10⁶ cells in 30 µL; SAP medium:Cultrex = 2:1) or subcutaneously (1 × 10⁶ cells in 100 µL; 4:1). Doxycycline (200 µg/mL) was provided in drinking water starting one day after recovery from surgery. Tumor growth was assessed by calipers (s.c. injection) or palpation (orthotopic injection) at regular intervals. Tumor volume was calculated as L×W²/2. Mice were monitored daily and euthanized when tumor volume exceeded 1 cm³ or when humane endpoints were reached (≥15% weight loss, impaired mobility).

### Cell Culture

#### iPSC Reprogramming and Culture

Skin biopsies from healthy donors (Ctrl7, Ctrl10, Ctrl14) and neuroblastoma patients (NB1, NB2, NB4) were reprogrammed as described previously^40,41,45,72^. iPSCs were maintained feeder-free in mTeSR™ Plus (STEMCELL Technologies, 100-0276) on tissue-culture–treated 6-well plates coated with Laminin-521 (BioLamina, LN521; 5%) at 37 °C, 5% CO₂. Cell cultures were tested for the presence of Mycoplasma routinely using the MycoAlert Mycoplasma Detection Kit (Lonza, LT07-318) following the manufacturer’s protocol.

For passaging, wells were rinsed with Ca²⁺/Mg²⁺-free DPBS (Sigma-Aldrich, D8537) and incubated with Gentle Cell Dissociation Reagent (STEMCELL, 100-0485) for 4–6 min at room temperature. Suspended colonies were collected in mTeSR Plus using a cell scraper, residual cells were recovered by rinsing with medium, and aggregates were re-plated at ∼50 clumps/cm².

### Differentiation to Trunk Neural Crest Cells (tNCCs) and Sympathoadrenal (SA) Lineage Cells

Differentiation from iPSCs to tNCCs and SA lineage cells was based on a combination of published protocols^37–39^ with minor adaptations. iPSCs were dissociated with TrypLE™ Select (Thermo Fisher A1217701) and plated at 25,000 cells/cm² on 1% Cultrex Stem Cell-Qualified RGF BME (R&D Systems 3434-010-02). Cells were cultured for 2 days in NCC#1 medium [DMEM/F-12 GlutaMAX Supplement (Thermo Fisher 31331093), N2 (Thermo Fisher 17502001), CHIR99021 (3 µM; Tocris 4423), and Y-27632 (10 µM; BD 562822)], followed by 4 days in NCC#2 [DMEM/F-12 GlutaMAX Supplement, N2, DMH1 (1.2 µM; Tocris 4126), and BMP4 (15 ng/mL; Thermo Fisher PHC9534 or Proteintech HZ-1045)]. tNCCs were isolated using EasySep™ Release Human PSC-Derived Neural Crest Positive Selection (STEMCELL 100-0047). For SA differentiation, tNCCs were plated at 100,000 cells/cm² on 1% Cultrex and cultured in SAP medium [BrainPhys™ Neuraonal Medium (STEMCELL 05790), B27 (Thermo Fisher 17504044), N2, GlutaMAX (Thermo Fisher 35050061), MEM NEAA (Thermo Fisher 11140050), SHH (50 ng/mL; R&D 1845-SH-100), Purmorphamine (1.5 µM; Merck SML0868), and BMP4 (50 ng/mL)]; Y-27632 was included for the first 24 h. From day 4, cells were switched to SAM medium [BrainPhys™ + B27, N2, GlutaMAX, MEM NEAA, BDNF (10 ng/mL; Thermo Fisher AF-450-02), GDNF (10 ng/mL; Thermo Fisher AF-450-10), and NGF (10 ng/mL; Thermo Fisher AF-450-01-B)] with medium changes every other day.

### Genetic constructs and nucleofection

To enable doxycycline-inducible MYCN expression in iPSC-derived late-stage cells, the *MYCN* ORF was cloned into a PiggyBac-based inducible backbone. PB-TRE3G-MYCN was a gift from David Vereide (Addgene plasmid #104542; http://n2t.net/addgene:104542; RRID: Addgene_104542)^73^ and XLone-eGFP NLS (no drug resistant gene) was a gift from Xiaoping Bao (Addgene plasmid #140032; http://n2t.net/addgene:140032; RRID: Addgene_140032). In-Fusion® cloning (Takara, 638947) generated XLone-MYCN_mCherryNLS (XM) by (i) replacing EF1α-EGFP with EF1α-mCherry-NLS in the XLone backbone and (ii) inserting MYCN from PB-TRE3G-MYCN using SpeI/KpnI (NEB) to replace EGFP.

XM together with CAG-hyPBase PiggyBac transposase (gift of Prof. Johan Holmberg, Umeå University), was co-nucleofected into iPSC lines using a Nucleofector™ IIb (Lonza) with program B-016. After two passages, mCherry⁺ cells were sorted on a Sony SH800S. Sorted derivatives were designated Ctrl10XM, Ctrl14XM, NB1XM, and NB4XM.

### Immunofluorescence

Cells on coverslips were fixed in 4% paraformaldehyde (PFA; Thermo Fisher, 28908) for 10 min at room temperature (RT) and washed twice with DPBS. Blocking was performed for 1 h at RT in DPBS + 0.05% Tween-20 with 5% donkey or goat serum (matched to secondary antibody host). Primary antibodies diluted in blocking buffer were incubated overnight at 4 °C in a humidified chamber. After returning to RT for 30 min, samples were washed twice with PBS-T (DPBS + 0.05% Tween-20), incubated with secondary antibodies, washed twice, stained with DAPI (0.02%; Sigma, D9542; 5 min, RT), and mounted in Dako Fluorescent Mounting Medium (Agilent, S3023). Primary/secondary antibodies: OCT4 (1:50, Santa Cruz sc-9081), NANOG (1:50, Santa Cruz sc-293121), SSEA4 (1:50, Santa Cruz sc-21704), SOX10 (1:250, CST 89356), HOXC9 (1:100, Abcam ab50839), PHOX2B (1:100, Santa Cruz sc-376997), PRPH (1:100, Millipore AB1530), Ki67 (1:250, Abcam ab16667), TH (1:100, Abcam ab112), TUBB3 (1:1000, BioLegend 801202), ISL1 (1:100, DSHB 39.4D5); donkey anti-mouse Alexa Fluor 488 (1:1000, Thermo Fisher A-21202), F(ab’)₂ goat anti-rabbit Alexa Fluor 568 (1:1000, Thermo Fisher A-21069).

Images were acquired on a ZEISS LSM 900-Airyscan at the Biomedicum Imaging Core (BIC), KI. For imaging, a 2×2 tile scan mode was applied, and stitched automatically using ZEN software to correct overlap and alignment.

### Immunohistochemistry

Formalin-fixed, paraffin-embedded (FFPE) tumor sections were processed by heat-induced antigen retrieval in 10 mM Tris-citrate (pH 6.0) or Tris-EDTA (pH 9.0) with 0.1% Tween-20 for 20 min in a pressure cooker, blocked in 10% donkey serum in TBS + 0.1% Tween-20, and subjected to Avidin/Biotin blocking (Vector, SP-2001). Primary antibodies (overnight, 4 °C): Ki67 (1:100, Abcam ab16667), PHOX2B (1:100, Abclonal A21990). The next day, endogenous peroxidase was quenched with 3% H₂O₂, and sections were incubated with biotinylated secondary antibody (Vector BA-1000; 2 h, RT), followed by ABC reagent (Vector PK-6100; 30 min). Detection used DAB, with hematoxylin counterstain, graded ethanol dehydration, and mounting. Slides were scanned on a ZEISS Axioscan 7 (20×) at BIC.

### Polymerase Chain Reaction and Sanger Sequencing

Genomic DNA was extracted and quantified by NanoDrop2000. Exon 25 of ALK was amplified using primers (Forward 5′-AATCCTAGTGATGGCCGTTG-3′; Reverse 5′-CCACACCCCATTCTTGAGG-3′) with DreamTaq DNA polymerase (Thermo Fisher, EP0703) under standard conditions (95 °C 1 min; 35 cycles of 95 °C 30 s, 58 °C 30 s, 72 °C 30 s; final 72 °C 10 min). Amplicons (∼200 bp) were verified on 1% agarose gels, purified, and subjected to Sanger sequencing at the KIGene core facility, KI. Data was analyzed using Snapgene.

### Quantitative Real-Time PCR (qRT-PCR)

Total RNA was extracted at indicated differentiation stages using Direct-zol™ DNA/RNA Miniprep (Zymo Research, R2052). cDNA synthesis used SuperScript™ III First-Strand Synthesis SuperMix (Thermo Fisher, 11752050) per manufacturer’s instructions. qPCR was performed with iTaq™ Universal SYBR® Green (Bio-Rad, 1725125) on a CFX384 Touch Real_Time PCR (Bio-Rad). Cq values were analyzed using the instrument software.

### Western Blot

Cells were lysed on plate in RIPA (Thermo Fisher 89900) supplemented with HALT™ protease/phosphatase inhibitor (Thermo Fisher 78442). After addition of LDS sample loading buffer (Thermo Fisher NP0007), 20µg of protein per lane was resolved on Bolt™ 4-12% Bis-Tris (Thermo Fisher NW04125BOX, Extend Figure 8b) or 8% SDS-PAGE (Figure 4c) gels. Proteins were transferred using Trans-Blot^®^ Turbo™ Mini Nitrocellulose (Extend Figure 8b, Bio-Rad, 1704158) or polyvinylidene difluoride (PVDF) membrane (Figure 4c, Millipore IPVH0010). Membranes were blocked in 5% BSA in PBST (0.05% Tween-20) for 1h, incubated overnight (4 °C) with primary antibodies—phospho-: pALK (CST 6941 1:2000), ALK (CST 3633, 1:2000), AKT (proteintech 10176-2-AP, 1:2000), phospho-AKT (CST 4060, 1:2000), ERK1/2 (BD Bioscience 610124, 1:3000), phospho-ERK1/2 (CST 4370, 1:2000), MYCN (Proteintech 10159-2-AP), GAPDH (Santa Cruz sc-32233, 1:1000 or sc-44724, 1:5000)—and then with HRP-conjugated goat anti-rabbit (Thermo Fisher 65-6120) or goat anti-mouse (Dako P0447) secondary antibodies.

### Flow Cytometry and Cell Cycle Analysis

Cell-cycle profiling was performed using the Click-iT™ EdU Alexa Fluor™ 647 Flow Cytometry Assay Kit (Thermo Fisher, C10635) according to the manufacturer’s protocol. Cells were incubated with EdU (10 µM) for either 4 h (tNCC stage) or 24 h (SAP stage) to account for differences in doubling time, then washed twice with PBS and fixed in 4% paraformaldehyde for 10 min. For ALK inhibition experiments, SAP-stage cells were treated with lorlatinib (PF-06463922, MedChemExpress HY-12215) or DMSO control 24 h before EdU incubation. EdU detection was performed following the kit protocol, and samples were passed through 100 µm strainers before acquisition on a BD FACSCanto II (BD Bioscience). Data were analyzed with FlowJo software (V10, BD Bioscience). For FACS of SAP_5, ERBB3-PE (clone 1B4C3, BioLegend 324705) and NGFR (D4B3, CST 8238S) were used; NGFR was conjugated to AF647 via FlexAble™ 2.0 Antibody Labeling Kit (Proteintech, KFA503). SAP cells from 3 independent inductions were stained and sorted on a Sony SH800S. Sorted SAP_5 and Negative fractions were replated at equal numbers in SAM medium. After 8 days, confluence was monitored every 4 h for 10 days using IncuCyte; plates were then processed for crystal violet staining.

### Bulk RNA Sequencing

#### Sample and Library Preparation

Total RNA (Direct-zol kit) was submitted to Novogene (UK) for quality control (QC), poly(A)-selected library preparation, and Illumina PE150 sequencing (∼6 GB raw data per sample). Reads underwent QC and adapter trimming, then were quantified using a quasi-mapping approach to transcripts in GRCh38.p13(Ensembl v109) using Salmon v1.9.0, and reported as TPM and counts. Upstream processing (QC, trimming, quantification) was performed on UPPMAX Rackham.

#### Quantification, Preprocessing, and Differential Expression

Transcript-level quantifications from Salmon were imported into R (v4.4.1)^74^ and summarized to the gene level using *tximport*^75^ with the transcript-to-gene mapping derived from Ensembl v109/GENCODE v43 (GRCh38.p13). Genes with fewer than five total counts across all samples were excluded, Gene IDs were harmonized with GENCODE v43, and sample metadata was curated before normalization. Gene-level counts were then analyzed with DESeq2^76^ using the design formula *∼CollectedTimePoint * Group*. Multiple testing was controlled by the Benjamini–Hochberg procedure (FDR < 0.05).

### Functional enrichment and visualization

GO and gene set enrichment analyses were performed with clusterProfiler (v4.10.1)^77^ and GSEA (v4.3.2)^78^. Plots summarizing expression dynamics across time points and groups were created with heatmaps (ComplexHeatmap^79,80^) and bubble/box plots (ggplot2^81^).

### Single-Cell RNA Sequencing

#### Library Generation and Sequencing

Single-cell suspensions thawed from frozen stocks were barcoded with TotalSeq™-B Antibody-Oligo conjugates (BioLegend B0251–B0254) per manufacturer workflow. Pooled samples were processed with the 10x Genomics Single Cell 3’ v3.1 workflow. Library construction, HTO enrichment, and indexing followed the TotalSeq workflow at the NGI-ESCG core facility (SciLifeLab, Stockholm). Libraries were sequenced on an MGI DNBSEQ PE100 flow cell (Xpress Genomics, Sweden).

#### Raw Data Processing and Quality Control

Raw sequencing data were processed with Cell Ranger v7.2.0 (10x Genomics) for sample-level de-multiplexing and alignment to the 10x Genomics reference bundle, generating count matrices per sample. Matrices were imported into R (v4.4.1)^74^ for downstream analysis using Bioconductor (v3.19) and Seurat (v4.4.0)^82^. Quality control was performed in R, removing outliers based on feature counts, UMI counts, and mitochondrial/ribosomal transcript percentages using median ± 5×MAD thresholds^83,84^. After filtering, 30,654 high-quality cells remained (NCC: 8,710; SAP: 9,680; D8: 5,761; D12: 6,503), which were then assembled into a Seurat object with *Read10X*.

### Data Integration, Normalization, and Clustering

*HTODemux* was used for HTO-based demultiplexing; singlets were retained for analysis. Datasets were normalized with *SCTransform*^85^ (mitochondrial %, ribosomal %, and cell-cycle scores regressed), and integrated across time points and conditions using Seurat’s standard integration workflow. The integrated object was then split into Ctrl-only and patient-derived (NB) subsets for comparative analyses. Dimensionality reduction was performed by PCA (top 50 PCs) and UMAP embeddings were computed from the top 22 PCs. Neighborhood graphs (*FindNeighbors,* top 22 PCs) and clustering (*FindClusters*) were constructed, with a clustering resolution of 0.2 chosen based on highest mean and minimum silhouette scores^86^. Marker genes for each cluster were identified using *FindAllMarkers* with Wilcoxon tests, applying Benjamini-Hochberg correction (FDR<0.05).

#### Transcriptional Signature Scoring and Cell Type Annotation

Sympathetic neuron (SYM) and mesenchymal (MES) markers were taken from Kastriti et al^44^ (Supplementary Data 2). Gene-module scores were further computed with *AddModuleScore* (Seurat). These scores were calculated as the difference between the average expression of the markers of interest (e.g. SYM or MES) and that of control genes matched for expression levels. A negative score indicates a higher expression of control genes relative to the markers of interest, while a positive score indicates the opposite. The SYM and MES scores for each cell positioned cells along a SYM–MES continuum. Cell-cycle phases were inferred using *CellCycleScoring* function from Seurat with default S-phase and G2/M-phase gene sets, which assigns each cell to G1, S, or G2/M phase. Cell-type annotation incorporated: time point, DEGs, cell-cycle state, pseudotime position (as explained next), cluster-specific programs, and SYM–MES scores.

The activity of the ALK pathway was estimated for each cell using as a proxy with the same scoring approach described above. In detail, the difference between the average expression of genes from the pathway annotated by Reactome (R-HSA-201556, also include in Supplementary Data 1) and that of control genes matched for expression levels were computed for each cell with *AddModuleScore*. Similarly, “SAP_5 module” scores were estimated by the same approach, using as reference the DEGs (log₂FC > 0.5, FDR < 0.05) derived from *FindAllMarkers*, for the SAP_5 cluster.

### Trajectory and Pseudotime Analysis

UMAP embeddings from Seurat were transferred to Monocle3^87^ (v1.3.1) and Slingshot^88^ (v2.7.0) for lineage reconstruction, and trajectories were visualized by cluster, sample, and pseudotime position. In Monocle3, dimensionality reduction was performed with 50 PCs (*num_dim = 50*), and pseudotime was assigned with the root node defined as NCC-1 (NCC-like cells) to reflect the expected developmental origin. In Slingshot, trajectories were inferred using Seurat clusters on the UMAP space (*clusterLabels = "cluster_ids", reduction = "umap"*), with NCC-1 set as the starting cluster.

### Cross-Projection of Patient-Derived and Ctrl Cells

To transfer labels between datasets, we implemented two complementary approaches. Using the SCP framework^89^, NB query cells were mapped to Ctrl reference cells with *RunKNNPredict* (k = 30, default), assigning each query to its nearest neighbors in the reference embedding. Predicted labels were inferred from majority neighbor votes, with confidence derived from neighbor vote distributions. Using Seurat utilities, we built transfer anchors between the integrated Ctrl reference and the combined dataset (*FindTransferAnchors*, top 22 PCs), and projected query cells onto the Ctrl UMAP manifold with *MapQuery* (reduction.model = "umap", seed = 22). Predicted cell classes were transferred from the reference collected time point to NB query cells. In both approaches, assignments were summarized by UMAP overlays for spatial context and by barplots quantifying cross-condition correspondences.

### Surface Marker Analysis

Cluster-specific DEGs for SAP_5 were obtained with *FindAllMarkers* (adj. P < 0.01; positive log₂FC). Genes were annotated via org.Hs.eg.db: GO:0005886 (plasma membrane) terms were retained. A gene-level specificity score was defined as

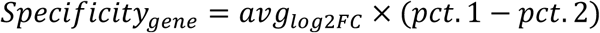

where *pct.1* and *pct.2* are detection fractions in SAP_5 versus all other clusters.

Top surface genes were evaluated pairwise. SCT-normalized expression > 0.5 denoted positivity. Double-positive fractions were computed in SAP_5 and other clusters:

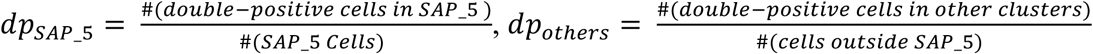

Pair specificity was defined as 𝑆𝑝𝑒𝑐𝑖𝑓𝑖𝑐𝑖𝑡𝑦*_pair_*_=_ = 𝑑𝑝*_SAP_5_* − 𝑑𝑝*_others_*.

A weighted specificity was computed as

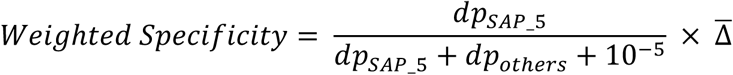

where Δ̄ is the mean log2 fold-change of the two genes. Gene pairs were ranked by weighted specificity, and top were selected for validation.

### Analysis of Patient Datasets in the R2 Online Database Tool

Analysis was performed in the online R2 platform^90^ on the datasets referred to in the text as SEQC^91^. Expression of SAP_5 marker genes and ALK activity-related genes was used to stratify patients into relatively high- and low-expression groups, either by KNN-based grouping or by median split following Cox proportional hazards regression. These stratifications were then used to perform Kaplan-Meier survival analysis, Cox regression modeling, correlation analysis between SAP_5 and ALK activity scores, and to assess the association of SAP_5 scores with MYCN amplification status and with age at diagnosis.

### Statistical analysis

Statistical analysis was performed in R (v4.4.1) or Prism (GraphPad v9). Data are presented as mean ± SD or mean ± SEM, from 6 independent biological replicates, as indicated in figure legends. Comparisons between two groups were performed using two-tailed Student’s *t*-tests, and comparisons among multiple groups were analyzed by one-way ANOVA followed by Šídák’s multiple-comparisons test. Chi-square tests were used for categorical variables, and the Wilcoxon rank-sum test was applied for single-cell data. Two-way ANOVA was used to assess time, group, and interaction effects in growth curve analyses. Multiple testing correction was performed using the Benjamini–Hochberg false discovery rate (FDR) method. Statistical significance was defined as *P* < 0.05 (adjusted *P* < 0.05 for multiple comparisons, as indicated).

## Data availability

Raw and processed single-cell RNA-seq and bulk RNA-seq data generated in this study will be deposited in the European Genome-phenome Archive (EGA). Publicly available single-cell RNA-seq datasets from hESC-based *in vitro* differentiation and normal human adrenal gland development used in this study are available under accession codes GSE219153 (Gene Expression Omnibus), GSE147821, and EGAS00001004388. Publicly available mouse NCC-derived lineage data are accessible under accession code GSE201257. Bulk RNA-seq data from neuroblastoma tumors were obtained from GSE62564.

## Code availability

All custom computer code used for data analysis in this study will be made available through our GitHub repository.

## Acknowledgement

We are grateful to all members of Wilhelm Lab (KI) for helpful discussions. We would like to acknowledge support from the National Genomics Infrastructure in Stockholm funded by Science for Life Laboratory, the Knut and Alice Wallenberg Foundation and the Swedish Research Council, and SNIC/Uppsala Multidisciplinary Center for Advanced Computational Science for assistance with massively parallel sequencing and access to the UPPMAX computational infrastructure. The computations and data handling were enabled by resources in projects [NAISS 2025/22-344, and NAISS 2025/23-207] provided by the National Academic Infrastructure for Supercomputing in Sweden (NAISS) at UPPMAX and PDC, funded by the Swedish Research Council through grant agreement no. 2022-06725. We would like to thank the core facilities at Karolinska Institutet: Biomedicum Flow Cytometry core facility, Biomedicum Imaging Core, KIGene, iPS Core facility, and Histology facility theme cancer. This work was supported by: Swedish Childhood Cancer Foundation (PR2021-0080, PR2024-0094, HFT-0001(MW), and TJ2021-0137, PR2021-0129 (O.C.B-R), Swedish Cancer Society (22-2236Pj, (MW), and 20-1159Pj (AF)), Swedish Research Council (2020-1427, 2023-02206 (MW), and 2022-00899 (AF)), Radiumhemmets Forskningsfonder (#214173 (MW)), KI Forskningsbidrag grant 2022-01925 (O.C.B-R), and Karolinska Institutet (2-5586/2017 (MW)).

---

**Extended Data Figure 1.**
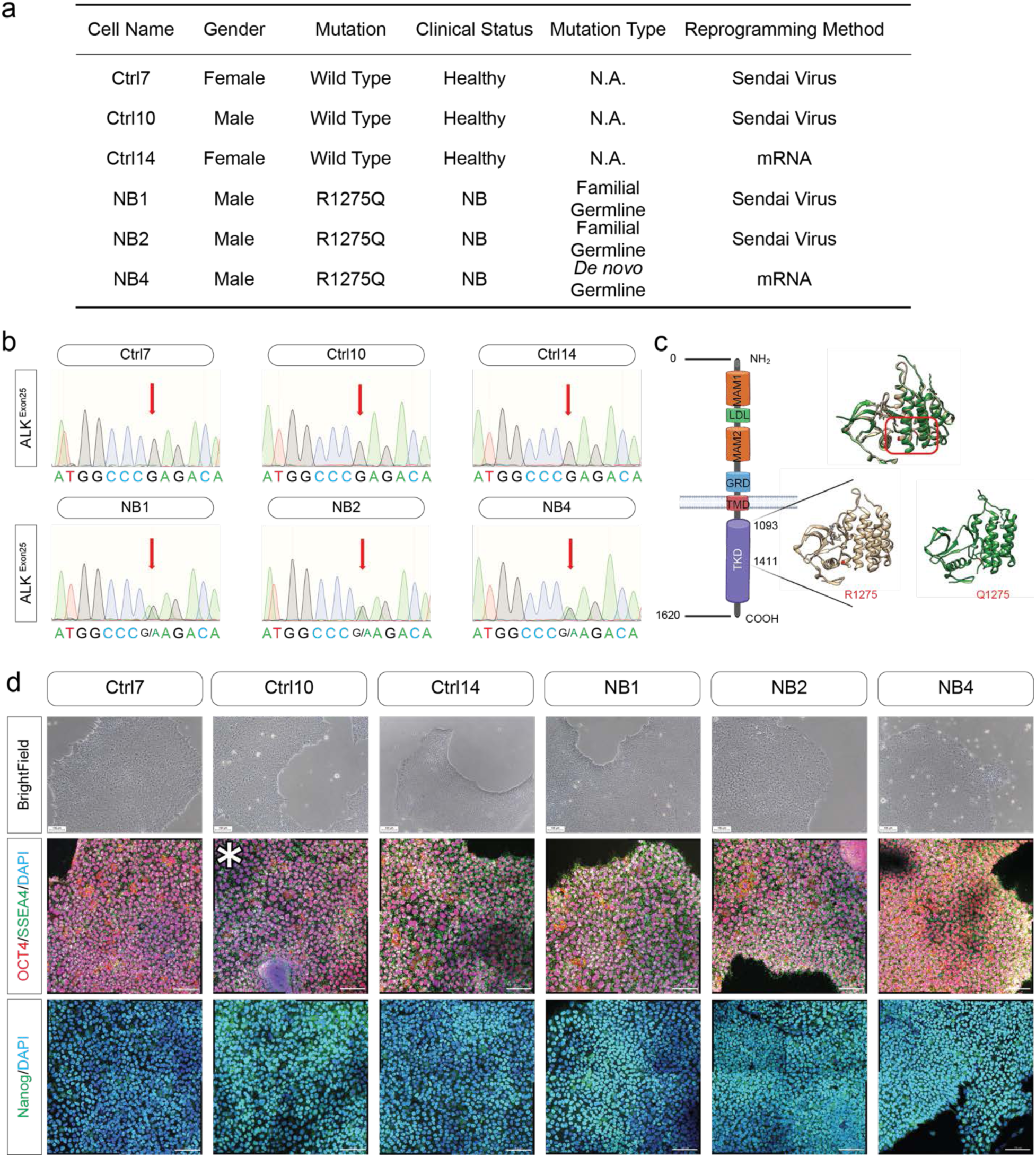
Sample information and iPSC reprogramming. **a,** Table summarizing donor information for all samples used in this study. **b,** Sanger sequencing of ALK exon 25 showing the germline mutation site. The red arrow indicates the mutated nucleotide. Nucleotide types are labeled below each peak; the mutation site is indicated as “G/A.” **c,** Schematic representation of ALK receptor structure and its functional domains (left). Protein structures of the tyrosine kinase domain (TKD) from wild-type (R1275, yellow, PDB ID: 5KZ0), mutant (Q1275, green, PDB ID: 4FNX), and the merged overlay (top) are shown on the right. **d,** Representative bright-field and immunofluorescence images showing iPSC morphology and expression of pluripotency markers OCT4, SSEA4, and NANOG. Scale bars: 100 µm. The image marked with an asterisk is also shown in Fig. 1a. The separated channels of the immunofluorescence are shown in Supplementary Fig. 1.

**Extended Data Figure 2.**
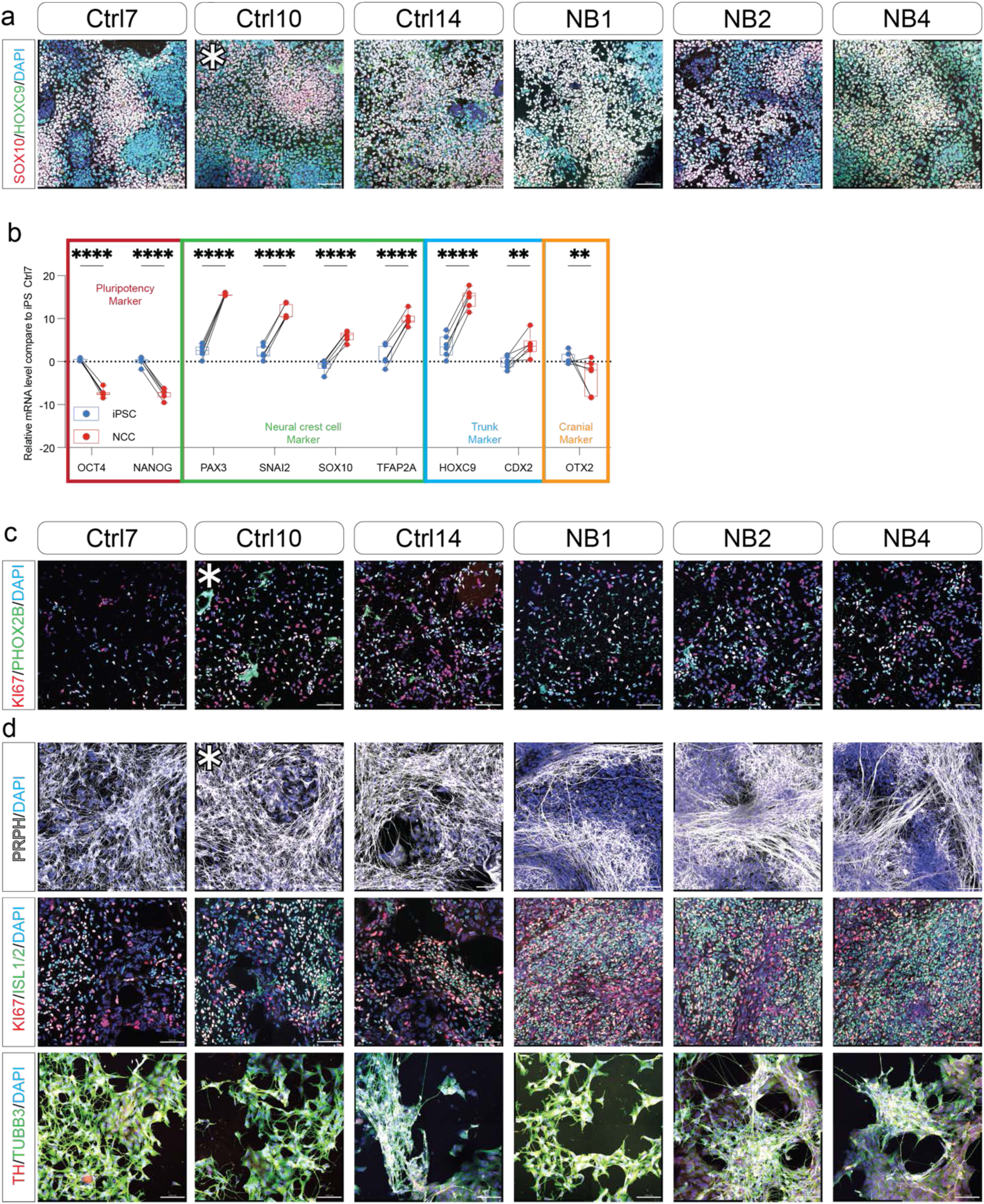
Validation of trunk neural crest cell (tNCC) and sympathoadrenal (SA) lineage cell differentiation. **a,** Representative immunofluorescence images showing tNCC expression of trunk marker HOXC9 and neural crest marker SOX10. Scale bars, 100 µm. The image marked with an asterisk is also shown in Fig. 1a. The separated channels of the immunofluorescence are shown in Supplementary Fig. 2. **b,** RT–qPCR analysis of marker expression across differentiation stages. Markers include pluripotency (*OCT4*, *NANOG*), neural crest (*PAX3*, *SNAI2*, *SOX10*, *TFAP2A*), trunk (*HOXC9*, *CDX2*), and cranial (*OTX2*). Each dot represents a biological replicate (n=6), and lines connect paired samples. Expression of each gene was normalized to Ctrl7 iPSCs. Data are shown as mean ± s.d. Statistical significance was determined using a paired two-tailed Student’s *t*-test. tNCC vs iPSC in *OCT4*, *NANOG*, *PAX3*, *SNAI2*, *SOX10*, *TFAP2A* and *HOXC9*, *P* < 0.0001 = ****; in *CDX2*, *P* = 0.002 = **; in *OTX2*, *P* = 0.0015 = **. **c,** Representative immunofluorescence images showing SAP expression of the sympathoadrenal lineage marker PHOX2B and the proliferation marker KI67. Scale bars, 100 µm. The image marked with an asterisk is also shown in Fig. 1a. The separated channels of the immunofluorescence are shown in Supplementary Fig. 3. **d,** Representative immunofluorescence images showing SAM expression of the peripheral nervous system (PNS) marker PRPH (top), the proliferation marker KI67, the mature sympathoadrenal marker ISL1/2 (middle), the catecholaminergic marker TH, and the neuronal marker TUBB3 (bottom). Scale bars, 100 µm. The image marked with an asterisk is also shown in Fig. 1a. The separated channels of the immunofluorescence are shown in Supplementary Fig. 4-5.

**Extended Data Figure 3.**
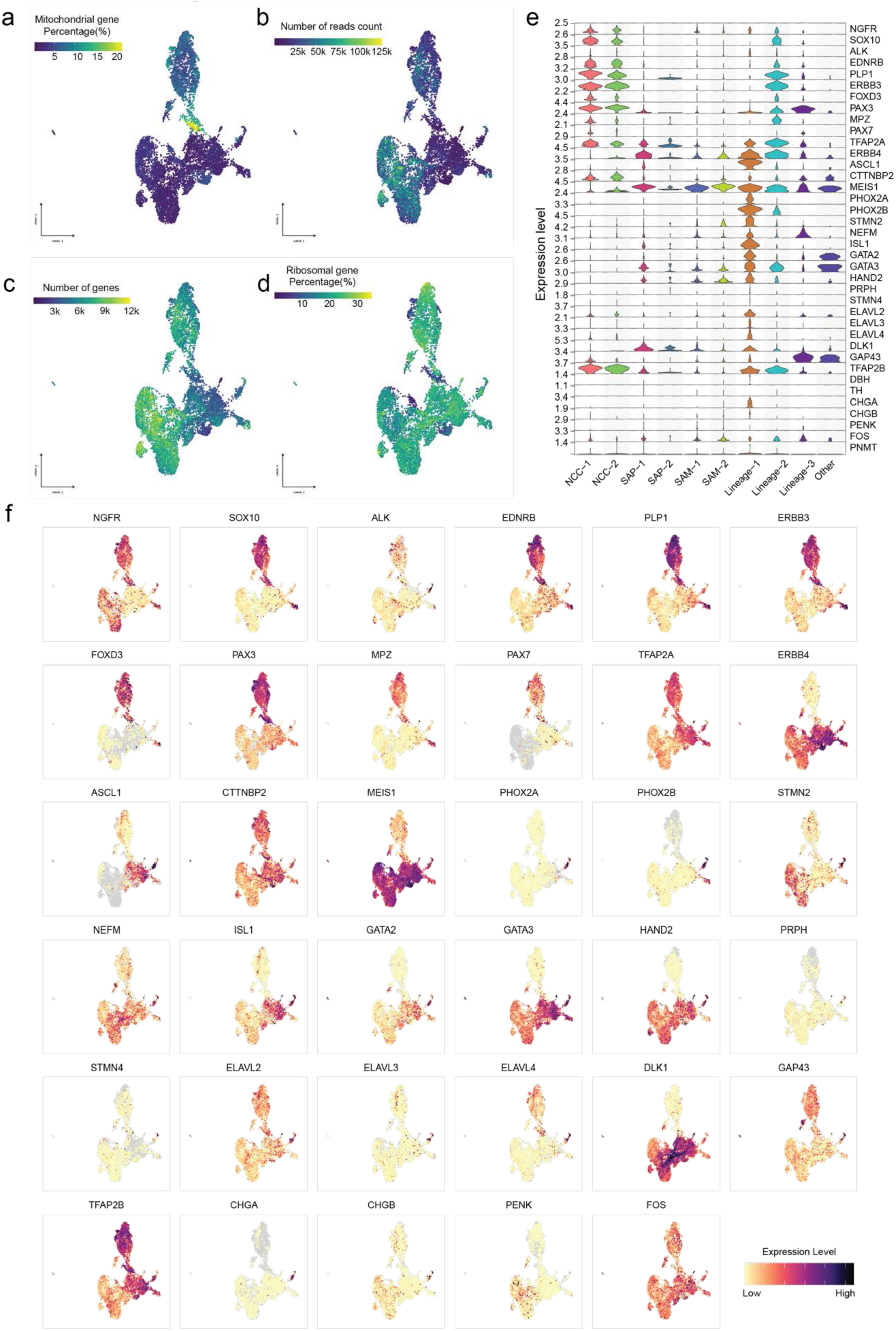
Quality control and marker gene visualization of single-cell RNAseq data from Ctrl cells. **a–d,** UMAP of Ctrl cells (as in Fig. 1d, middle) after quality-control filtering, colored by percentage of mitochondrial genes (a), number of reads per cell (b), number of detected genes (c), and percentage of ribosomal genes (d). **e,** Violin plots showing the expression levels of selected marker genes from published single-cell data of human adrenal medulla development^42,43^. **f,** UMAP of Ctrl cells (as in Fig. 1d, middle) colored by selected feature genes (same as in Supplementary Fig. 1e), with high expression in dark purple, low expression in yellow, and no expression in grey.

**Extended Data Figure 4.**
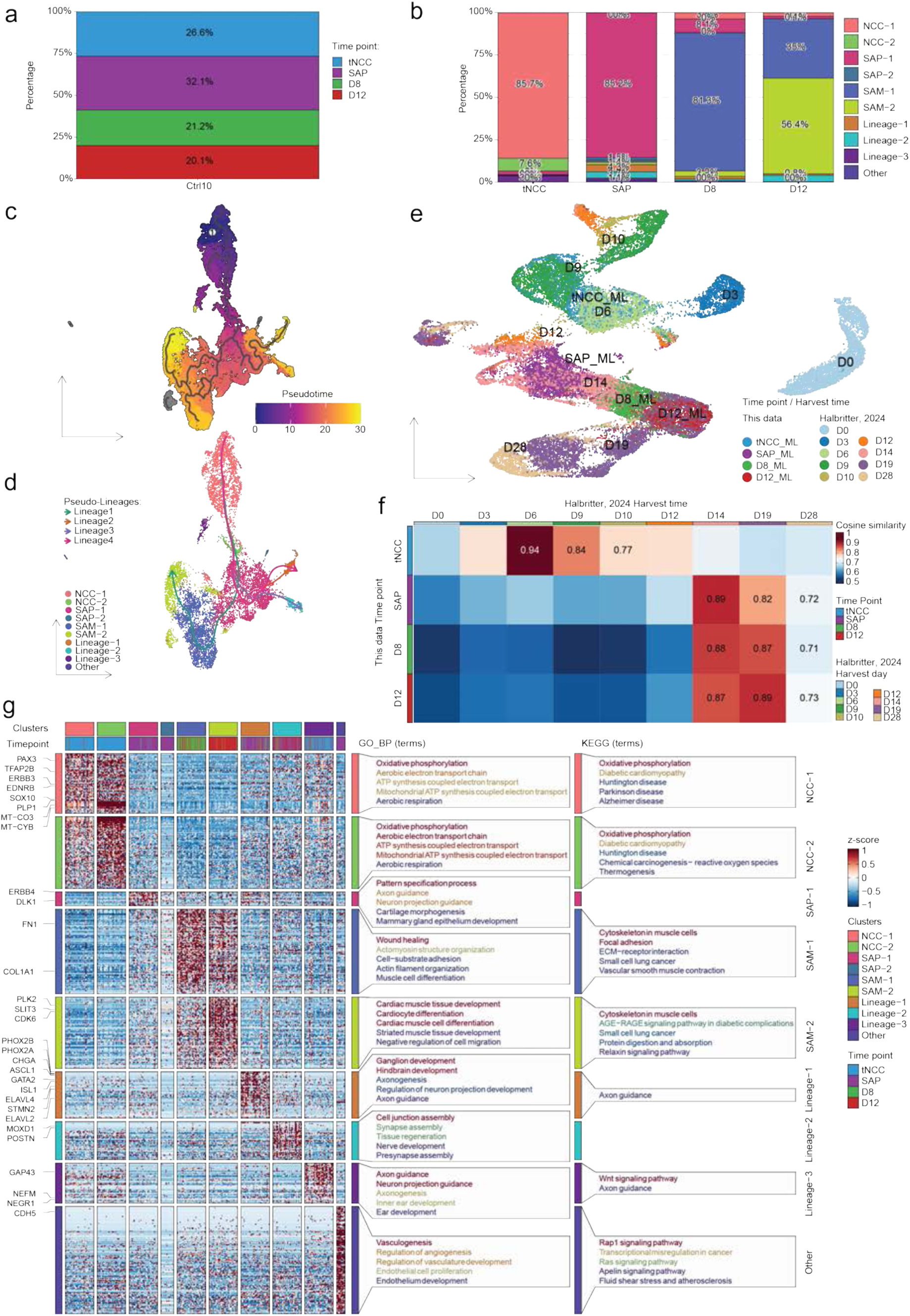
Cell distribution, trajectory inference and comparative analysis of single-cell RNAseq data. **a,** Proportion of cells from each collection time point in Ctrl lines. **b,** Proportion of clusters represented at each collection time point in Ctrl lines. **c,** Pseudotime analysis of Ctrl cells using Monocle3. UMAP (as in Fig. 1d, middle) is shown, with Monocle3-defined clusters in distinct colors. Lines indicate predicted developmental trajectories. Pseudotime is colored from dark purple (early) to yellow (late). The NCC-1 cluster was chosen as the root for trajectory inference. **d,** Pseudotime trajectory analysis of Ctrl cells using Slingshot. Arrows indicate predicted developmental trajectories. UMAP as in Fig. 1d, middle. **e,** Integrated UMAP of data from this study with a previously published hESC-based *in vitro* differentiation dataset^26^, colored either by collection time point (this study, same colors as in Fig. 1e, upper left) or by harvest time. **f,** Heatmap of cosine similarity between this study (query) and the reference dataset from Saldana-Guerrero et. al^26^. Scores are colored from red (high similarity, maximum = 1) to blue (low similarity, minimum = 0.5). Colors correspond to time points (this study) or harvest time (reference) as in Supplementary Fig. 6e. **g,** Heatmap of z-scored expression values of DEGs upregulated in each Ctrl cluster. Color scale represents high (red) to low (blue) expression. Genes associated with SA lineage differentiation are labeled on the left. GO biological process (GO_BP) and KEGG enrichment analyses of DEGs are shown on the right, with the top five terms (if applicable) displayed. Color denotes FDR values (red to blue).

**Extended Data Figure 5.**
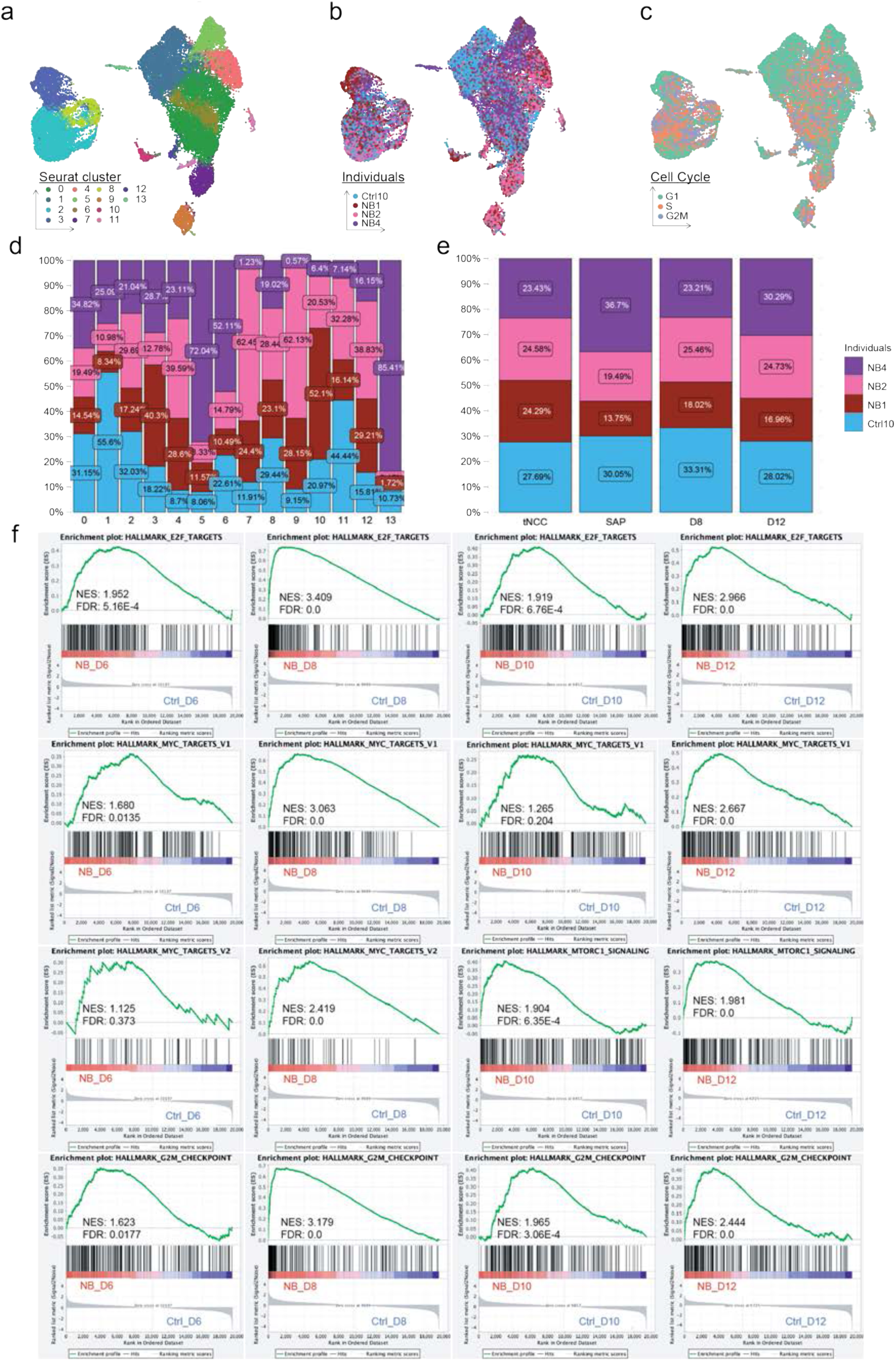
Overview of single-cell RNAseq data from all cells, including Ctrl and patient-derived lines, and GSEA of SA lineage. **a–c,** UMAP of all cells (as in Fig. 2b), colored by Seurat clusters (a), individual identity (b), and cell-cycle status (c). **d–e,** Proportion plots showing: individual identity by Seurat cluster (d) or by collection time point (e). **f,** Gene set enrichment analysis (GSEA) plots of proliferation-related pathways across SA lineage time points, showing pathways upregulated in patient-derived cells.

**Extended Data Figure 6.**
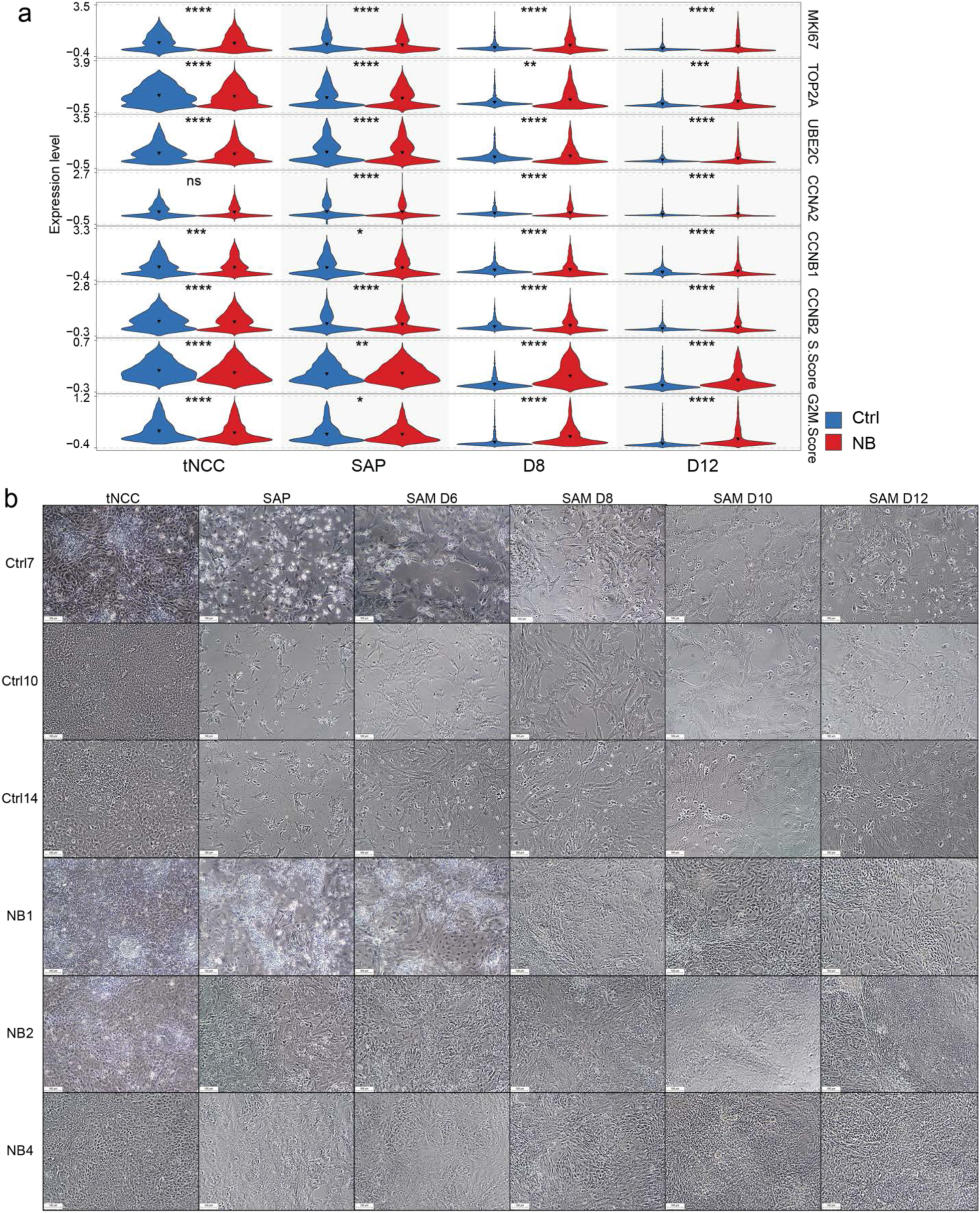
Patient-derived cells show increased proliferation compared to Ctrl cells during SA lineage commitment. **a,** Violin plots showing expression levels of selected proliferation-associated genes and cell-cycle phase scores (S phase and G2/M phase). *P* < 0.0001 = ****. In TOP2A, Ctrl vs NB at D8, *P* = 0.0012 = **; at D12 *P* = 0.0003 = ***. In CCNB1, Ctrl vs NB at SAP, *P* = 0.002 = *. In S score, Ctrl vs NB at SAP, *P* < 0.001 = **. In G2M score, *P* = 0.048 = *. **b,** Representative bright-field images at key time points prior to sample collection for sequencing, illustrating cell morphology and cell density. Scale bar, 100 µm.

**Extended Data Figure 7.**
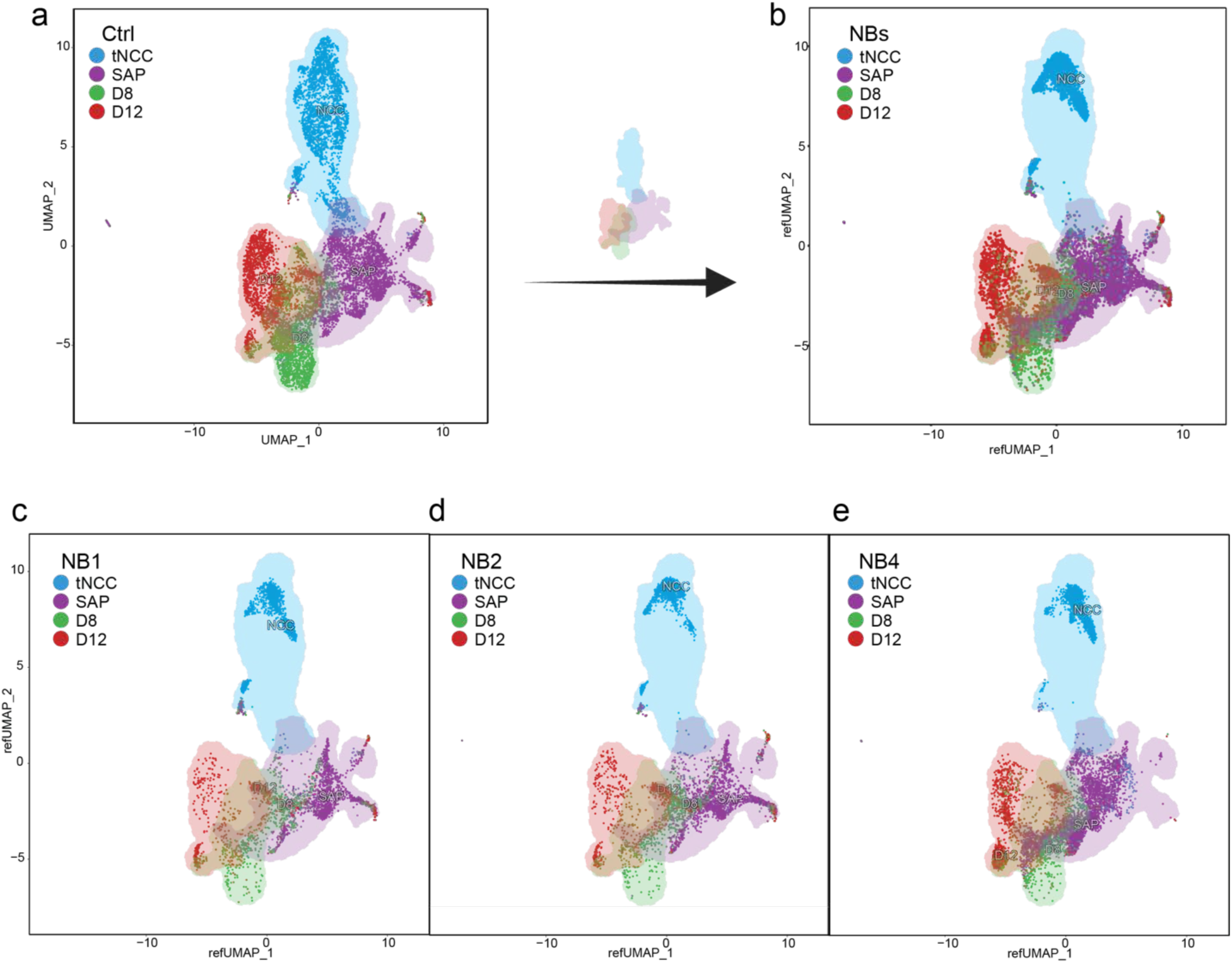
Mapping patient-derived cells onto the reference UMAP derived from Ctrl cells. **a,** Reference UMAP of Ctrl cells (as in Fig. 1d, middle), with shaded regions indicating different collection time points. **b,** All patient-derived cells were mapped onto the Ctrl UMAP using the *MapQuery* function in Seurat. **c–e,** Patient-derived cells from NB1 (c), NB2 (d), and NB4 (e) mapped onto the Ctrl UMAP.

**Extended Data Figure 8.**
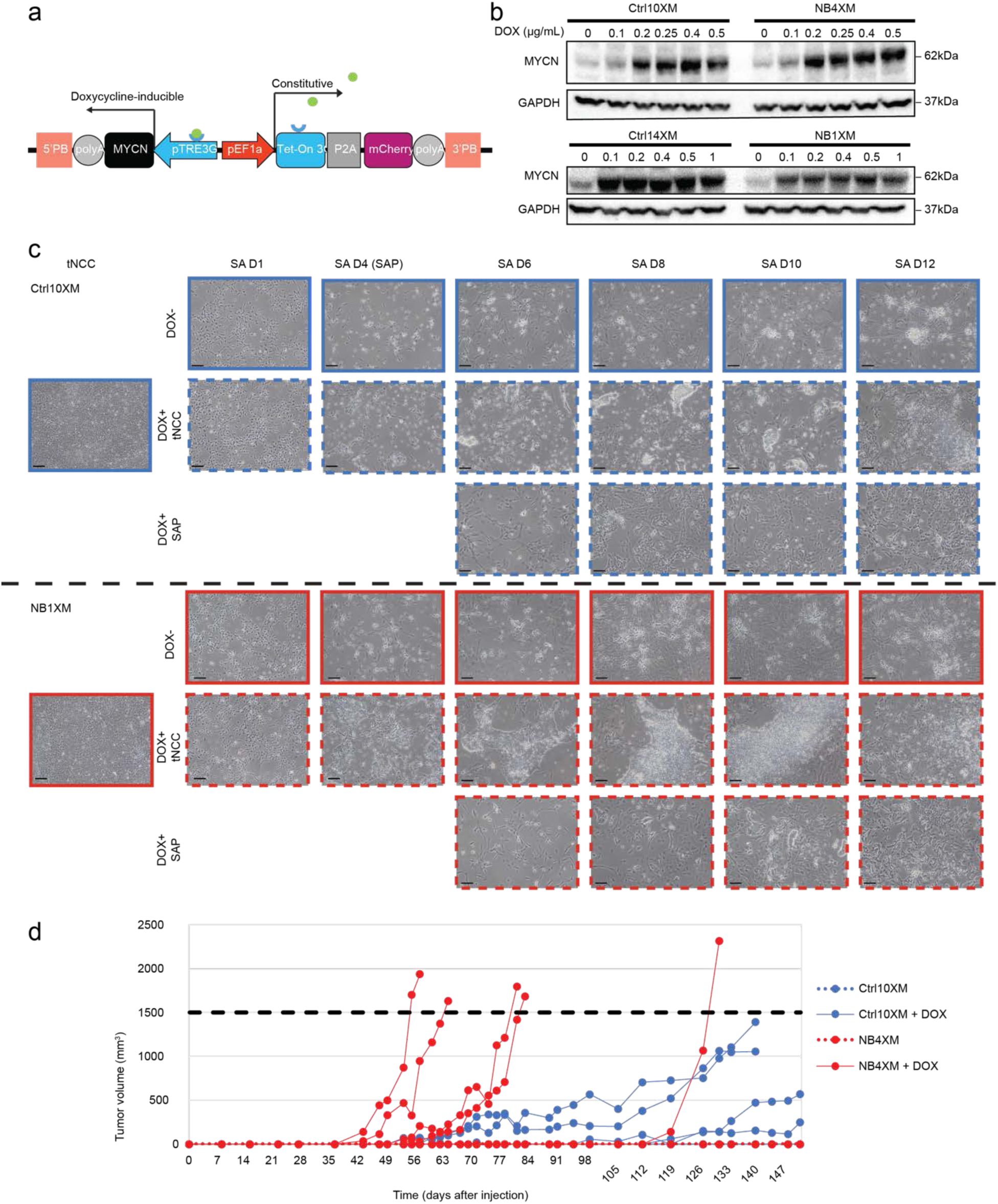
Generation of iPSCs with an inducible MYCN expression system and tumor growth following subcutaneous transplantation of tNCC in mice. **a,** Schematic of the integrated construct. The PiggyBac transposase integrates sequences between two PB arms (orange). The EF1α promoter (red) drives constitutive expression of Tet-On 3G (blue square, co-activator of the TRE3G promoter) and mCherry (cherry square), separated by a P2A peptide (grey square). Upon addition of doxycycline (green dots), Tet-On 3G activates the TRE3G promoter, which drives expression of the *MYCN* to produce its coding sequence (CDS, black square). **b,** Western blot validation of inducible MYCN protein expression in iPSCs treated for 24h with increasing concentrations of doxycycline. **c,** Bright-field images showing differentiation of iPSCs carrying the construct into the SA lineage via the tNCC stage. Differences were observed when doxycycline was added at the tNCC stage (middle row) or SAP stage (bottom row) compared with untreated controls, in both Ctrl (top) and patient-derived lines (bottom). Scale bar, 100µm. **d,** Tumor growth curves from the subcutaneous injection model. Each line represents an individual mouse. Solid lines indicate mice treated with doxycycline, dashed lines indicate untreated mice. The thick black dashed line marks the experimental endpoint at tumor volume 1,500 mm³.

**Extended Data Figure 9.**
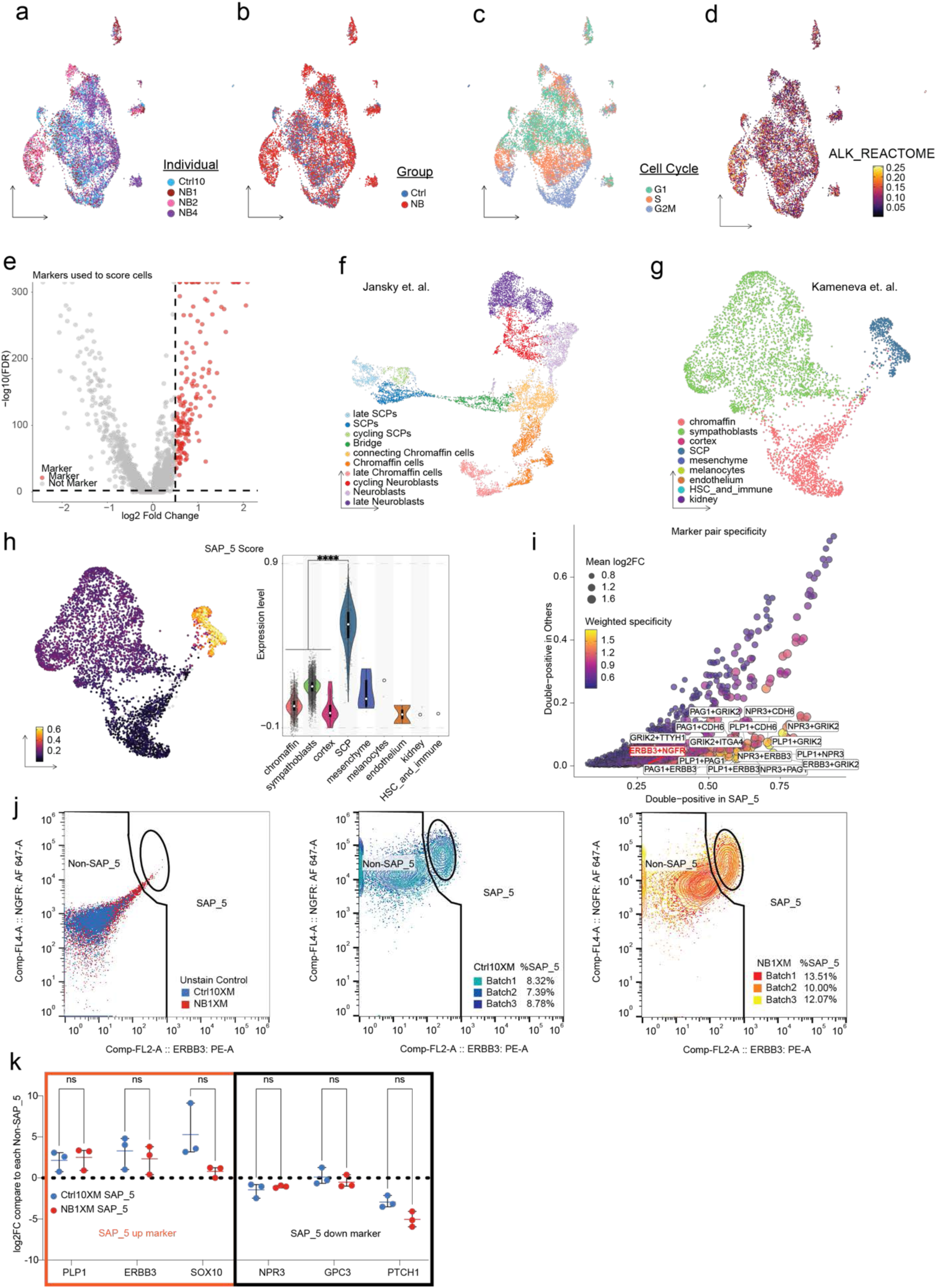
Characterization of SAP-stage cells and identification of SAP_5-specific gene expression. **a–d,** UMAP of SAP-stage cells (as in Fig. 5b), colored by individual identity (a), group (Ctrl versus NB; b), cell-cycle status (c), and ALK reactome activity score expression (d). **e,** Volcano plot showing DEGs in the SAP_5 cluster compared to all other SAP clusters. Red dots represent genes selected for module score calculation (log₂FC > 0.5 and adjusted P < 0.05). **f–g,** UMAP of published human adrenal medulla single-cell data^43^ colored by cell type annotations from the original study. Panel f shows the UMAP as in Fig. 5i (left). **h,** UMAP of the same published dataset (as in Extended Data Fig. 9g), colored by SAP_5 module scores (left). Violin plots show the distribution of SAP_5 scores across annotated clusters. Statistical significance was assessed using the Wilcoxon test. **i,** Dot plot showing marker-pair specificity in SAP_5 cells. Surface markers with high and differentially expressed levels were paired two at a time, and specificity scores were calculated for each pair. The x-axis shows the double-positive ratio in SAP_5 cells, and the y-axis shows the ratio in other clusters. Dot size represents the log₂FC of paired markers, and color represents weighted specificity (yellow, high; dark purple, low). Top-ranked marker pairs by weighted specificity are labeled in black. The validated surface marker pair used for downstream live-cell FACS sorting is labeled in red. **j**, Final SAP_5 gating across three independent inductions in both Ctrl (middle) and patient (right) lines, together with unstained control (left). **k**, RT–qPCR validation of selected marker expression by log_2_ fold change in sorted SAP_5 cells compared to their respective negative populations. Data are shown as mean ± s.d. The thick black dashed line marks log_2_FC = 0. Statistical significance was assessed using a two-tailed Student’s *t*-test.

**Extended Data Figure 10.**
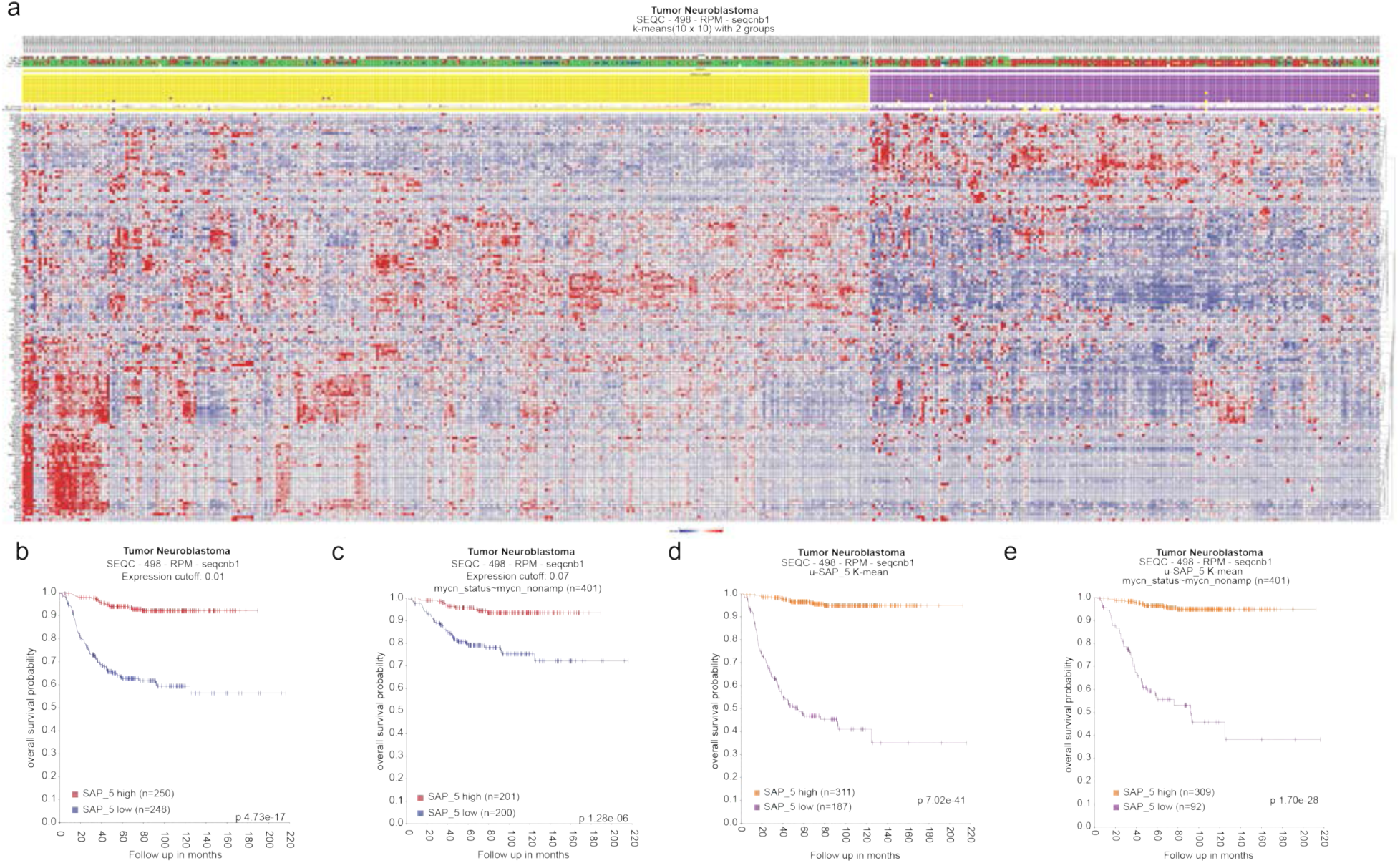
Association between SAP_5 marker expression and clinical outcomes in neuroblastoma. **a,** Heatmap of k-means clustering of SAP_5 cell markers. Each column represents an individual patient, and each row represents a SAP_5 marker. Patients were clustered into SAP_5-high (n = 311, orange) and SAP_5-low (n = 187, purple) groups. Marker expression values are shown as z-scores. **b–c,** Kaplan–Meier survival curves stratified by SAP_5 scores (median split: high versus low). Analyses were performed in all patients regardless of MYCN status (b, 498 patients in total, 248 patients in SAP_5 low group and 250 patients in SAP_5 high group) or restricted to MYCN non-amplified cases (c, 401 patients in total, 200 patients in SAP_5 low group and 201 group in SAP_5 high group). **d–e,** Kaplan–Meier survival curves stratified by k-means two-cluster grouping of SAP_5 markers, including all patients (d, 498 patients in total, 187 in SAP_5 low group and 311 in SAP_5 high group) or restricted to MYCN non-amplified cases (e, 401 patients in total, 92 in SAP_5 low group and 309 in SAP_5 high group).

**Supplementary Figure 1.**
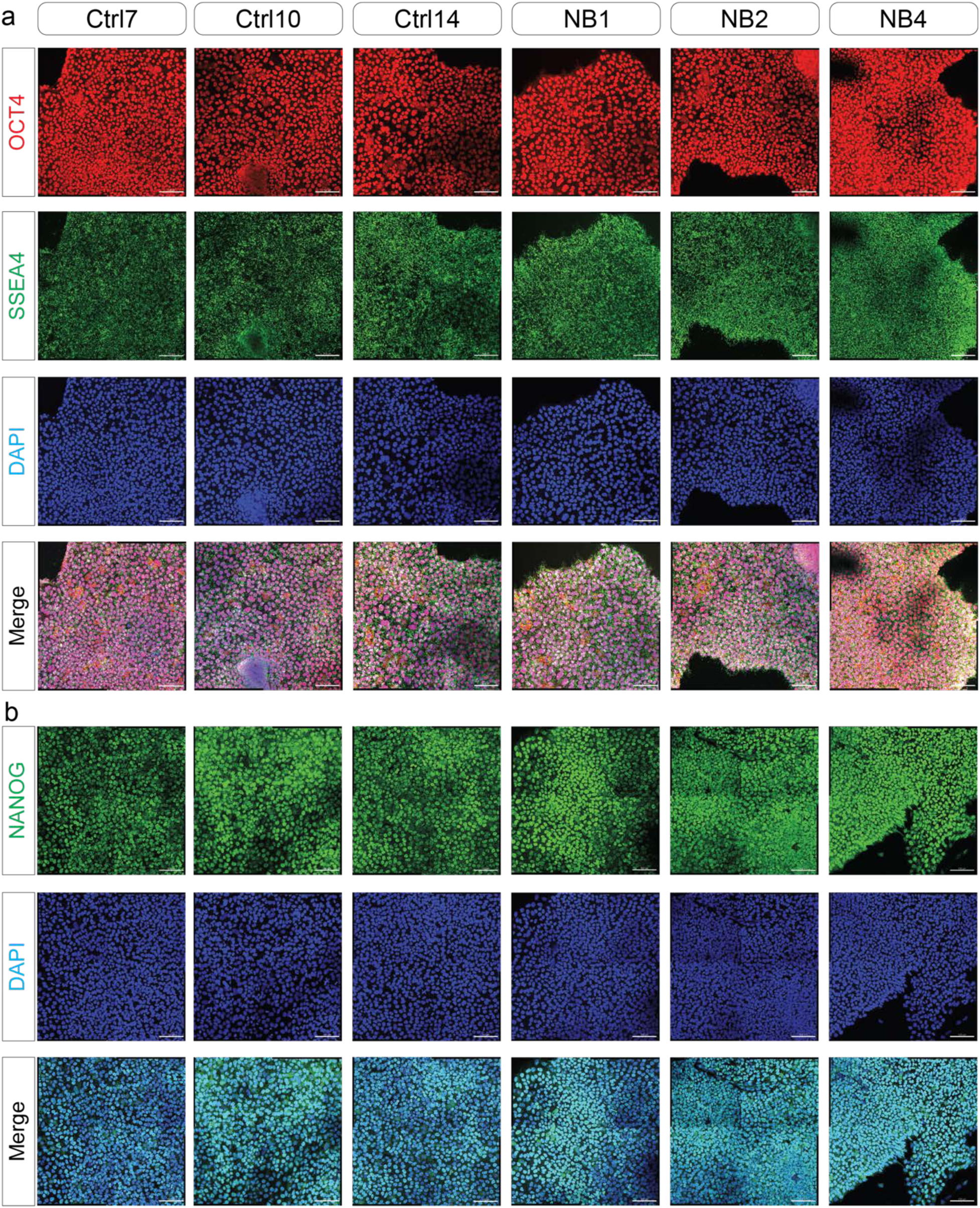
The individual channel of immunofluorescence of the iPSC stage. **a-b**, Representative immunofluorescence images on separated channels showing iPSC expression of pluripotent markers OCT4, SSEA4, and NANOG. DAPI stains the nuclei. Scale bars, 100 µm.

**Supplementary Figure 2.**
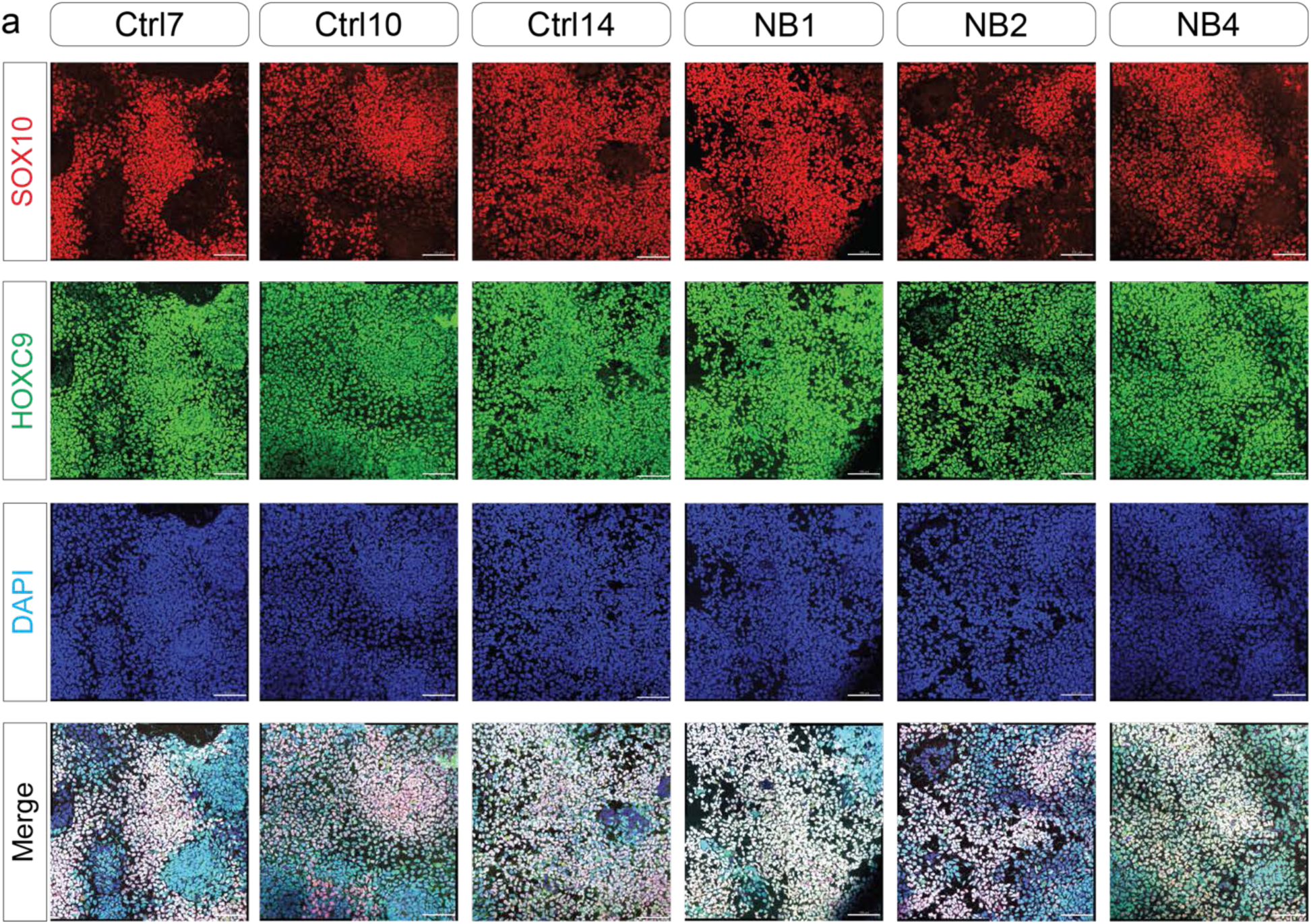
The individual channel of immunofluorescence of the tNCC stage. Representative immunofluorescence images on separated channels showing tNCC expression of neural crest marker SOX10, and trunk marker HOXC9. DAPI stains the nuclei. Scale bars, 100 µm.

**Supplementary Figure 3.**
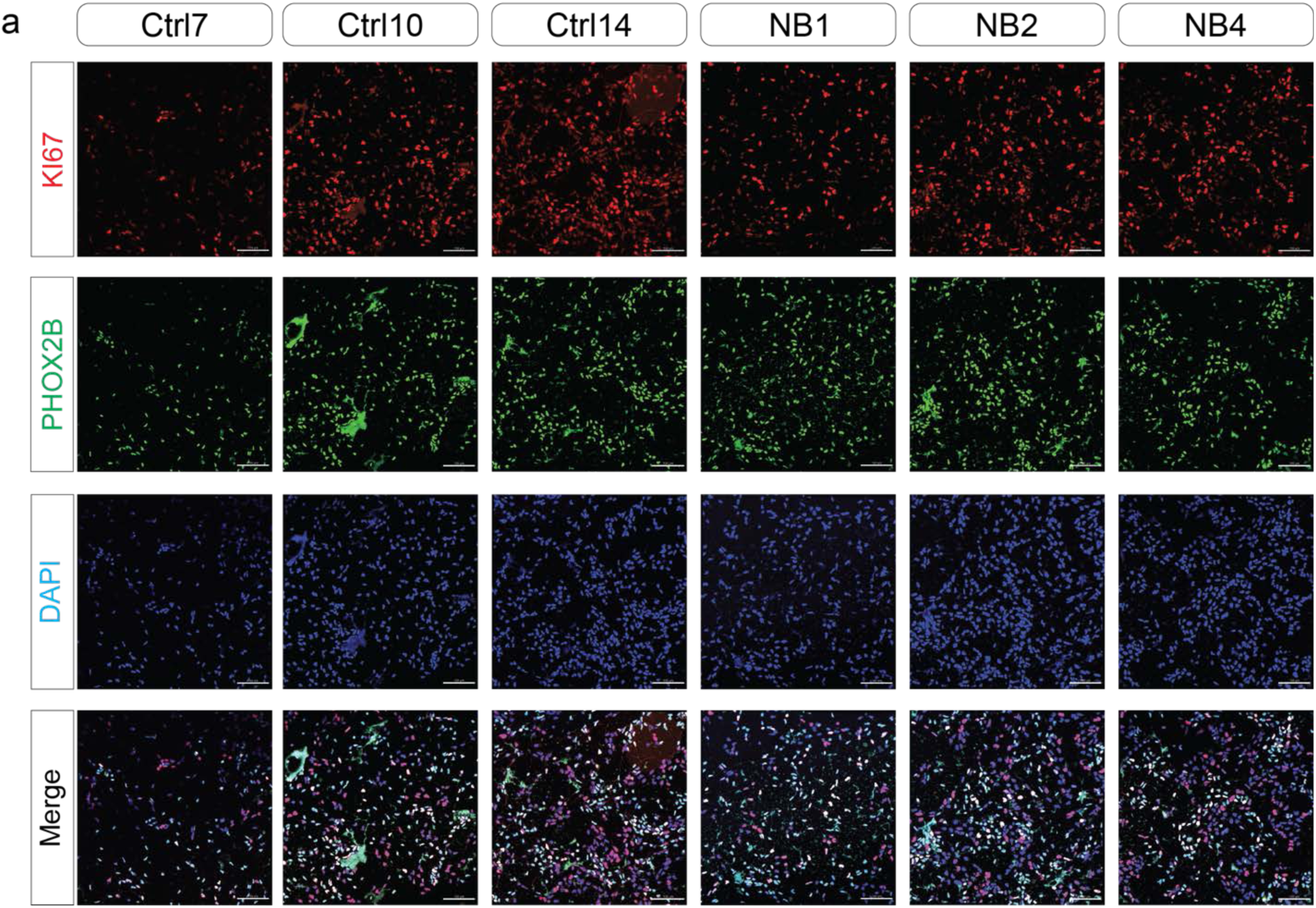
The individual channel of immunofluorescence of the SAP stage. Representative immunofluorescence images on separated channels showing SAP expression of proliferative marker KI67, and sympathoadrenal lineage marker PHOX2B. DAPI stains the nuclei. Scale bars, 100 µm.

**Supplementary Figure 4.**
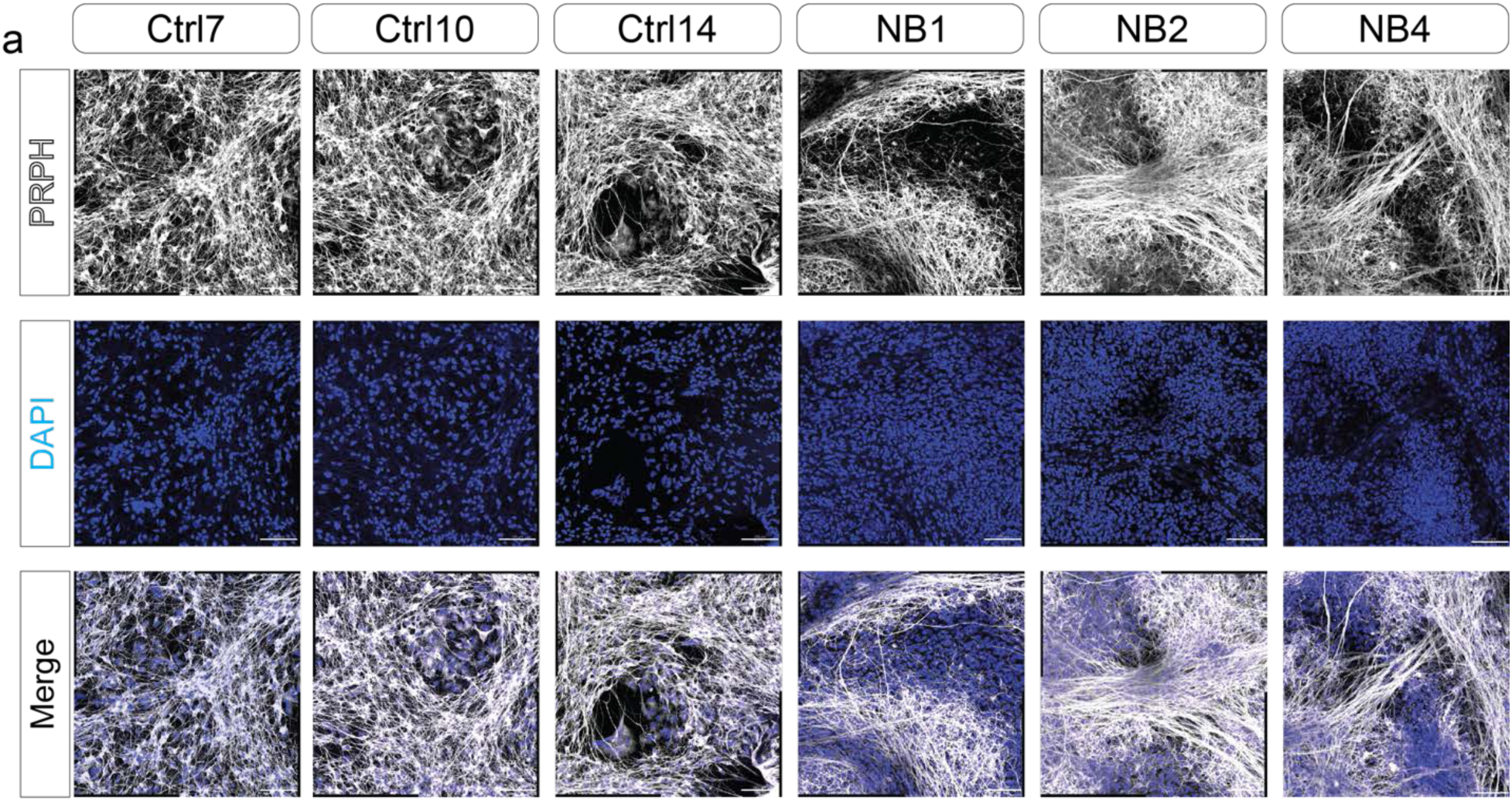
The individual channel of immunofluorescence of the SAM stage of PRPH. Representative immunofluorescence images on separated channels showing SAM expression of PNS marker PRPH. DAPI stains the nuclei. Scale bars, 100 µm.

**Supplementary Figure 5.**
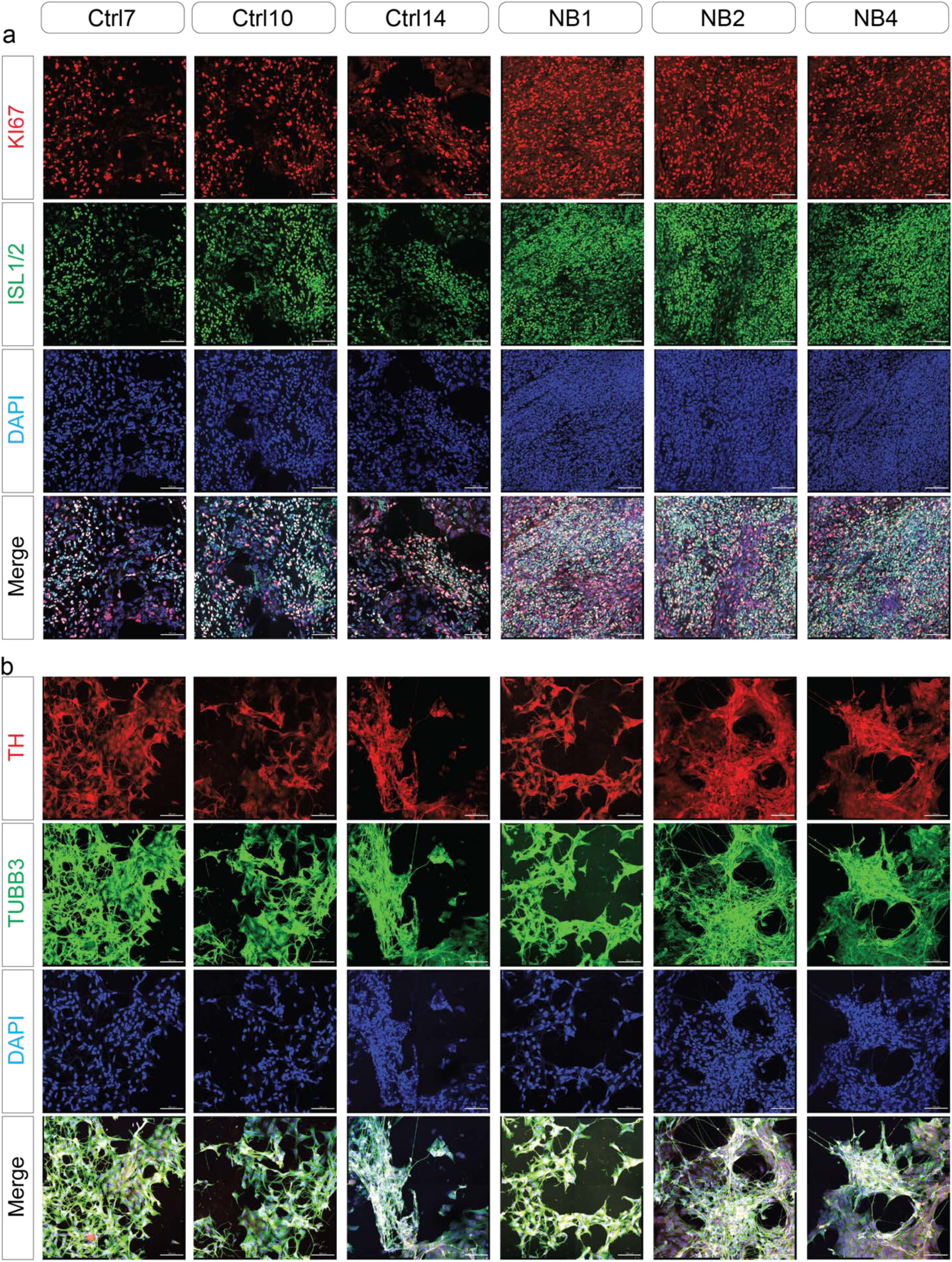
The individual channel of immunofluorescence of the SAM stage. **a-b**, Representative immunofluorescence images on separated channels showing SAP expression of the proliferation marker KI67, the mature sympathoadrenal marker ISL1/2 (a), the catecholaminergic marker TH, and the neuronal marker TUBB3 (b). DAPI stains the nuclei. Scale bars, 100 µm.

**Supplementary Figure 6.**
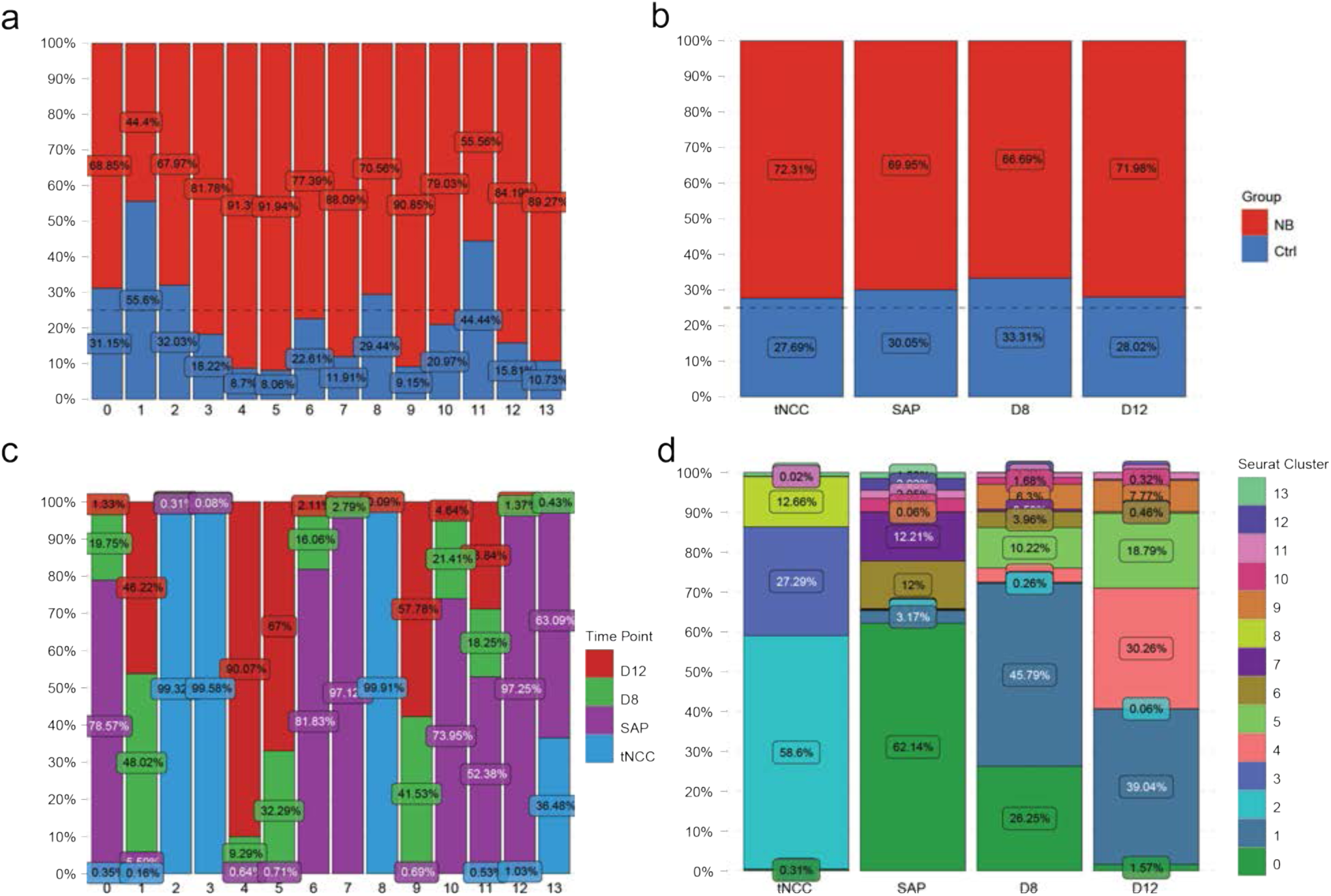
Overview of single-cell RNAseq data from all cells, including Ctrl and patient-derived lines. **a-d,** Proportion plots showing: group (patient versus Ctrl) by Seurat cluster (a) or by collection time point (b); collection time point by Seurat cluster (c); and Seurat cluster by collection time point (d).

**Supplementary Figure 7.**
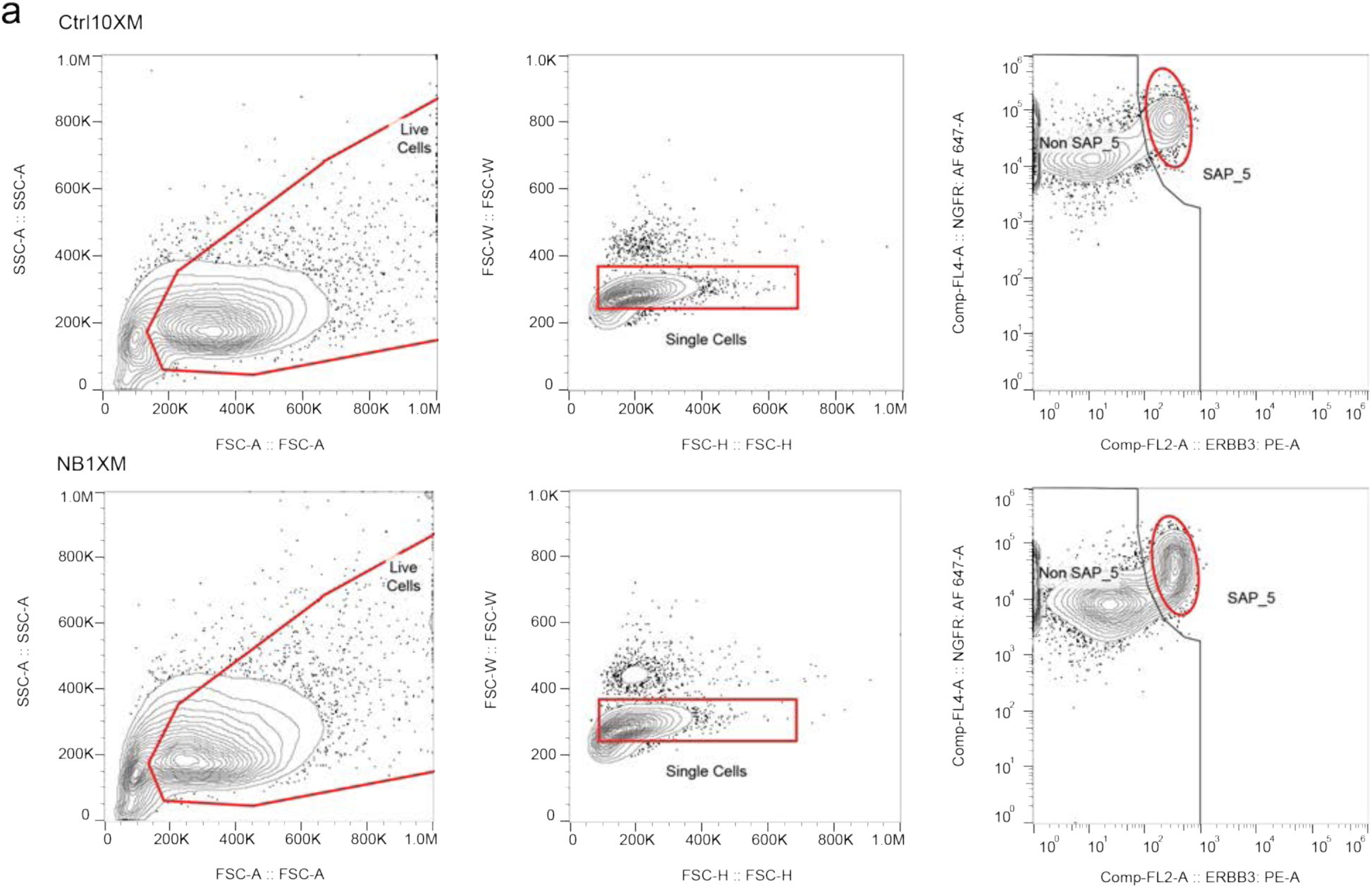
Gating strategy of SAP_5 cell sorting. Gating strategy for isolating SAP_5 cells from all events. The red region indicates cells selected for the next analysis or final SAP_5 gating.

